# Inhibition of mTOR decreases insoluble protein burden by reducing translation in *C. elegans*

**DOI:** 10.1101/2020.08.17.253757

**Authors:** Zhuangli Yee, Shaun Hsien Yang Lim, Li Fang Ng, Jan Gruber

## Abstract

Aging animals accumulate insoluble protein as a consequence of a decline of proteostatic maintenance with age. In *Caenorhabditis elegans*, for instance, levels of detergent-insoluble protein increase with age. In longer-lived strains of *C. elegans*, this accumulation occurs more slowly, implying a link to lifespan determination. We further explored this link, and found that detergent-insoluble protein accumulates more rapidly at higher temperatures, a condition where lifespan is short. We employed a *C. elegans* strain carrying a GFP transcriptional reporter under the control of a heat shock (*hsp-16*.*2*) promoter to investigate the dynamics of proteostatic failure in individual nematodes. We found that early, sporadic activation of *hsp-16*.*2* was predictive of shorter remaining lifespan in individual nematodes. Exposure to rapamycin, resulting in reduced mTOR signaling, delayed spurious expression, extended lifespan, and delayed accumulation of insoluble protein, suggesting that targets downstream of the mTOR pathway regulate the accumulation of insoluble protein. We specifically explored ribosomal S6 kinase (*rsks-1*) as one such candidate and found that RNAi against *rsks-1* also resulted in less age-dependent accumulation of insoluble protein and extended lifespan. Our results demonstrate that inhibition of protein translation *via* reduced mTOR signaling resulted in slower accumulation of insoluble protein, delayed proteostatic crisis and extended lifespan in *C. elegans*.

## 1. Introduction

Aging is marked by a gradual loss of homeostatic capacity, resulting in declining efficiency in the removal and repair of macromolecular damage and the eventual accumulation of such damage (Ogrodnik, Salmonowicz, & Gladyshev, 2019). Age-dependent failure of proteostasis is of particular interest, not least due to evidence that oxidatively damaged as well as insoluble protein accumulate with age, both in simple model organisms and vertebrates, including humans (Chiti & Dobson, 2017; Hipp, Kasturi, & Hartl, 2019). Accumulation of modified, misfolded, and insoluble protein has been suggested as a cause of aging in the nematode *Caenorhabditis elegans (C. elegans)* (David et al., 2010; Hartl, 2016; Kaushik & Cuervo, 2015; D. K. Kim, Kim, & Lee, 2016) and is known to play a causative role in several important age-related diseases in humans (David, 2012; Hartl, 2016; Klaips, Jayaraj, & Hartl, 2018; Polling, Hill, & Hatters, 2012). Failure of proteostatic processes and accumulation of insoluble protein is therefore an example of a “public” mechanism of aging (Kaushik & Cuervo, 2015; Partridge & Gems, 2002), suggesting that pathways and mechanisms affecting proteostasis may be conserved determinants of lifespan and health span. Consistent with this hypothesis, several known aging pathways have been shown to directly impact protein translation, proteome maintenance and the rate of age-dependent decline in proteostasis (Anisimova, Alexandrov, Makarova, Gladyshev, & Dmitriev, 2018; Campisi et al., 2019; Labbadia & Morimoto, 2014).

Reduced insulin/IGF-1 signaling (IIS), for example, increases longevity (Kenyon, Chang, Gensch, Rudner, & Tabtiang, 1993) and has been reported to decrease age-dependent protein aggregation in *C. elegans* (Cohen, Bieschke, Perciavalle, Kelly, & Dillin, 2006; David et al., 2010). Similarly, loss of the germline extends lifespan in nematodes, and this intervention which also reduces protein aggregation has been reported in both gonad- and germline-less worms (Andux & Ellis, 2008; David et al., 2010; Hsin & Kenyon, 1999; Khodakarami, Saez, Mels, & Vilchez, 2015; Maklakov & Immler, 2016). Importantly, in germline-competent but not in germline-deficient nematodes, key heat shock responses are blunted after animals reach maturity, indicating that the maturation of germline cell redirects resources earlier in life and uses those to maintain proteostasis (Labbadia & Morimoto, 2015; Shemesh, Shai, & Ben-Zvi, 2013).

Another mechanism known to modulate lifespan across large evolutionary distances is reduced signaling through the mTOR (mechanistic target of rapamycin) pathway (Kaeberlein, Burtner, & Kennedy, 2007). Rapamycin, the eponymous inhibitor of mTOR, extends lifespan in diverse organisms, including yeast, nematodes, flies and mice (Bjedov et al., 2010; Harrison et al., 2009; Powers, Kaeberlein, Caldwell, Kennedy, & Fields, 2006; Robida-Stubbs et al., 2012). Key processes controlled through mTOR include mRNA translation, metabolism, and protein turnover (Saxton & Sabatini, 2017). mTOR inhibition is further known to modulate proteostasis by increasing autophagy, predominantly through activation of the FoxA/*pha-4* transcription factor (Sheaffer, Updike, & Mango, 2008). Another downstream target of mTOR and rapamycin is the p70 ribosomal S6 kinase (S6K), with inhibition of mTOR signaling resulting in reduced protein translation (Saxton & Sabatini, 2017). This mechanism likely contributes to lifespan extension by mTOR inhibition, because the reduction in the abundance of the *C. elegans* homolog of S6K (*rsks-1*) alone is sufficient to extend lifespan (Hansen et al., 2007; McQuary et al., 2016; Pan et al., 2007). Reduced transcription and deletion mutation of S6K/*rsks-1* have been reported to extend the mean lifespan of *C. elegans* by between 9% and 20% (Chen et al., 2013; Pan et al., 2007). These observations suggest a direct link between mTOR-dependent lifespan benefits and reduced protein translation; an exciting possibility, given that reduced translation has been reported to improve proteostasis and extends lifespan in *C. elegans* even when initiated late in life (Solis et al., 2018). Interestingly, rapamycin is also similarly known to significantly extend lifespan in mice, even when administered late in life (Harrison et al., 2009). Previous reports have shown that mTOR inhibition using rapamycin activates protein degradation *via* the ubiquitin-proteasome system or autophagy, thereby enhancing the clearance of misfolded proteins, (Zhao, Zhai, Gygi, & Goldberg, 2015). This also reduces protein synthesis, another mechanism that may promote proteostasis (King et al., 2008). However, whether mTOR inhibition or loss of *rsks-1* reduces accumulation of detergent-insoluble protein *in C. elegans* has not been directly determined yet. Here, we therefore explore the hypothesis that key benefits of mTOR inhibition are mediated by inactivation of S6K/*rsks-1* and that this results in reduced detergent-insoluble protein and promotes proteostasis in *C. elegans*. We compared the effect of mTOR inhibition (rapamycin) and ablation (RNAi knockdown) of *rsks-1* on lifespan and the accumulation of detergent-insoluble protein and aging in *C. elegans*.

Lifespan and biomarker studies observing changes in cohorts of aging animals only provide insights regarding cohort-level averages of these parameters. This is an important limitation because there is significant heterogeneity in individual aging trajectories and age-dependent dysfunction of individual animals after exposure to stressors such as heat shock in cohorts of isogenic nematodes. Prior studies have demonstrated, for instance, that stochastic variation of *hsp-16*.*2* expression following heat shock treatment may predict subsequent variation in survival, and that this is the result of general differences in protein dosage (Burnaevskiy et al., 2019; Herndon et al., 2002; A. Mendenhall, Crane, Tedesco, Johnson, & Brent, 2017; A. R. Mendenhall et al., 2012; Rea, Wu, Cypser, Vaupel, & Johnson, 2005). Whether similar variations in survival may be observed in the absence of a stressor (without heat shock) remains to be explored, especially given that in a typical wild-type (WT) cohort maintained at the standard temperature of 20 °C, the first animal may die on Day 10 of life. In contrast, the most long-lived individual of the same cohort may survive beyond Day 30 (Herndon et al., 2002).

We asked whether aging trajectories and remaining lifespans of individual animals were related to the status of their individual proteostatic system. Apart from some strains designed to express specific, aggregation-prone proteins (Morley, Brignull, Weyers, & Morimoto, 2002), it is challenging to observe insoluble protein in individual animals directly. To address this problem and to directly investigate proteostatic failure during the aging of individual animals, we utilized a transgenic strain of *C. elegans* that carries a green fluorescent protein (GFP) reporter-gene construct under the control of the *hsp-16*.*2* (heat shock protein) promoter (Link, Cypser, Johnson, & Johnson, 1999). *Hsp-16*.*2* is a member of the small heat shock proteins (∼16 kDa) and is typically only expressed upon heat shock, during which it facilitates protein refolding (Haslbeck & Vierling, 2015). When determined following heat shock, the robust induction of this transcriptional reporter has been reported to associate with a longer lifespan in *C. elegans* (Rea et al., 2005).

Here, we were not interested in the canonical function of *hsp-16*.*2* following heat shock or its direct impact on lifespan but spurious activation of this pathway as a biomarker. Small heat shock proteins are components of the proteostatic stress responses and are therefore expected to be induced under conditions where insoluble protein burden or damage is increased significantly (Haslbeck & Vierling, 2015). Here, we hypothesized that in the absence of a stressor such as a heat shock, age-dependent, spurious induction of *hsp-16*.*2* signals a spontaneous crisis and collapse of the proteostatic system.

A fraction of animals in aging cohorts of *hsp-16*.*2p*::GFP animals show a significant age-dependent activation of this reporter, even in the absence of heat shock. We first quantified the age-dependent dynamics of this effect and then sorted aging *hsp-16*.*2p*::GFP animals based on threshold presence of such spurious GFP expression (in the absence of heat shock) (Fig. 2**a**). Finally, we determined if such spontaneous GFP expression was indicative of increased insoluble protein levels and if it was predictive of the remaining individual lifespan animals in the same cohort. Using this approach, we found that the spontaneous activation of the *hsp-16*.*2p*::GFP reporter indicates an acute, age-dependent collapse of the proteostatic system, suggesting that the proteostatic failure is an endogenous event in *C. elegans*, leading to a crisis state. This possibly is akin to age-dependent diseases in humans, which are also driven by proteostatic failure and aggregation of endogenous proteins. Finally, we investigated the effect of mTOR inhibition on this marker of proteostatic failure.

**Figure 1.**
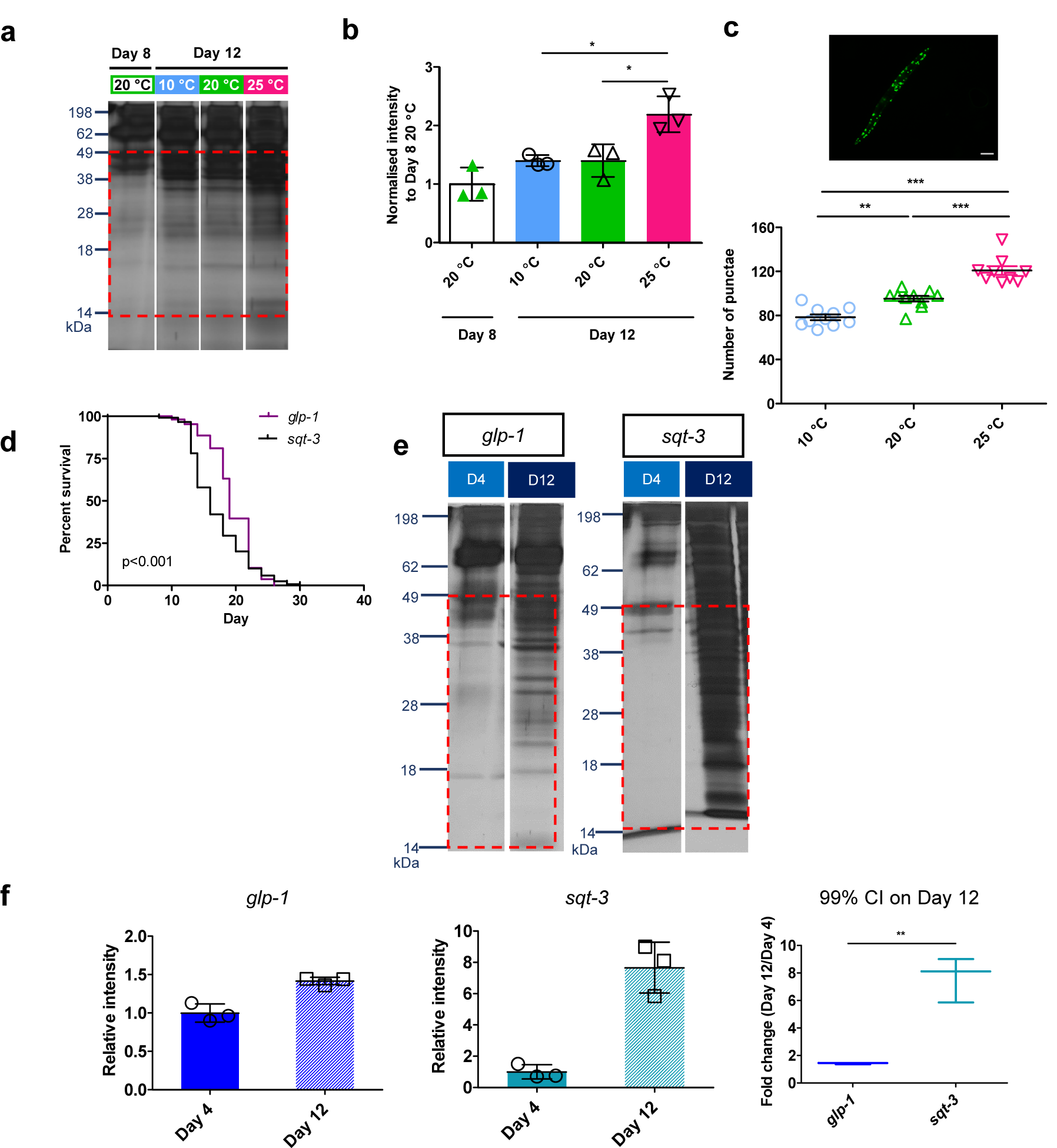
Accumulation of detergent-insoluble protein occurred more rapidly with higher temperatures and with functional germline in *C. elegans*. (a) For insoluble protein measurement assay, animals were age-synchronized and maintained at 20 °C until Day 8. One batch of animals was harvested as the reference on Day 8, before transferring the remaining cohort to fresh plates kept at either 10, 20 or 25 °C. (b) There was a general trend towards increased detergent-insoluble protein between Day 8 and Day 12 for all cohorts. The fold-increase (mean±SD) for all temperatures on Day 12, normalized to Day 8 were: 10 °C (1.4±0.1), 20 °C (1.4±0.3), and 25 °C, (2.2±0.3). Comparison of the normalized detergent-insoluble protein for 10, 20, and 25 °C on Day 12 showed that the means were significantly different (p<0.05, one-way ANOVA). (c) Representative image of an AM141 mutant animal (*unc-54p*::Q40::YFP) showing abundant polyQ punctae that can be used for quantification of protein aggregates in this strain. Scale bar, 100 µm. Mutants of *unc-54p*::Q40::YFP were raised at 20 °C before being transferred to 10, 20, and 25 °C on Day 8 of life. The number of intestinal polyQ punctae were quantified on Day 15 of life using an operator blinded scoring procedure. The abundance of punctae increased with temperature (p<0.001 for all comparisons, one-way ANOVA). Comparisons between groups are indicated in the figure. Data represented as Mean±SEM. (d) Comparison of survival of animals of the *glp-1* and *sqt-3* strains, maintained at 25 °C from Day 4 of life, was different between strains, with *glp-1* surviving significantly longer (p<0.001, Log-rank test). (e) Gel images showing the insoluble protein of three independent cohorts of animals. (f) Comparing fold changes between Day 4 and Day 12 between *glp-1* and *sqt-3* across all three independent experiments. The average fold change for *glp-1* was 1.4±0.05 (Mean±SD) while for *sqt-3*, it was 7.7±1.6 between Day 4 and Day 12. 99% confidence interval for these fold changes for *glp-1* and *sqt-3* do not overlap, suggesting that *sqt-3* accumulate abnormal protein significantly faster than *glp-1* (p<0.01, t-test).

**Figure 2.**
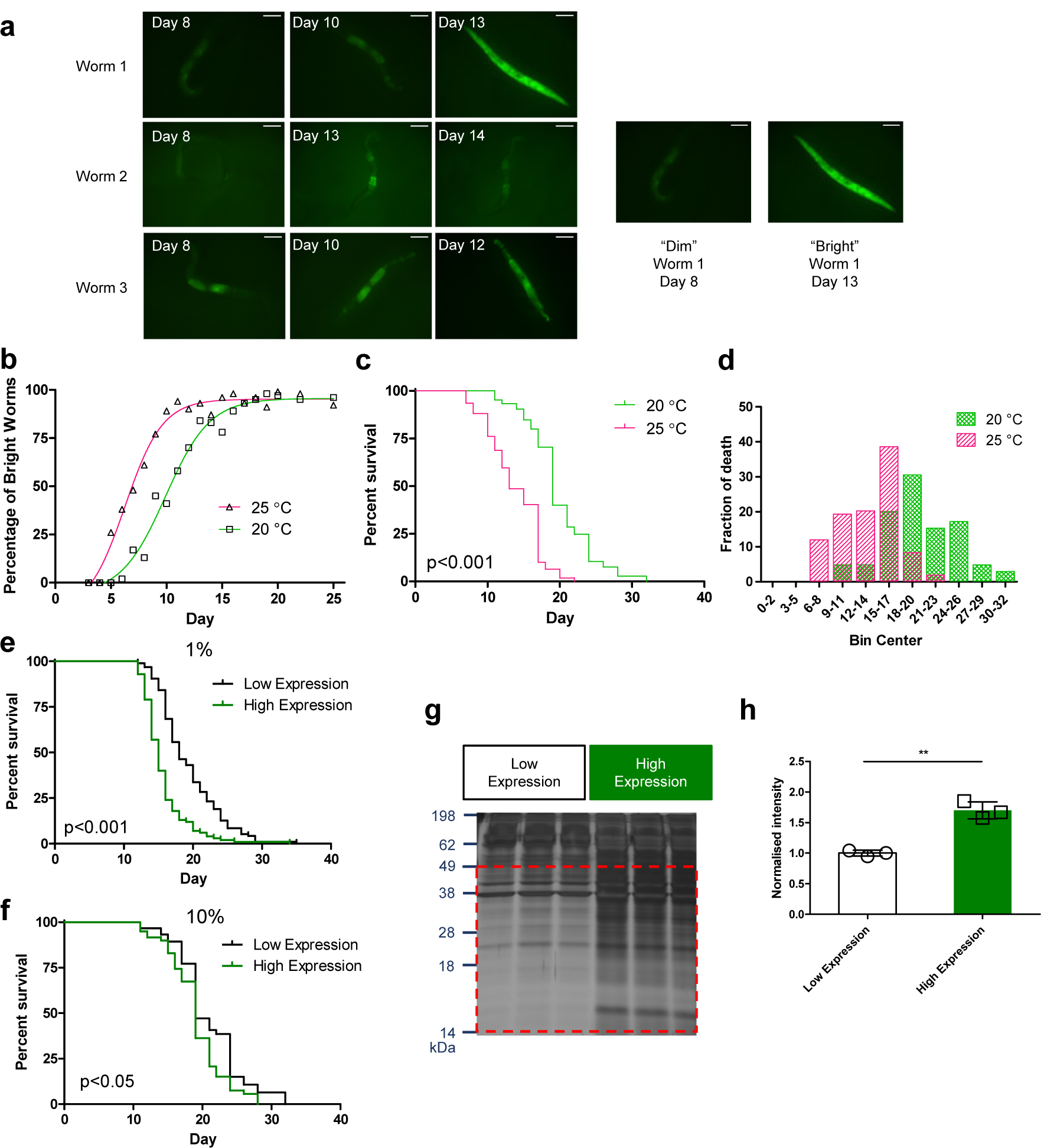
Stochastic *hsp-16*.*2p::*GFP expression correlates with a shorter lifespan. (a) The inter-individual variability in the age-dependent dynamics of *hsp-16*.*2p::*GFP expression (“dim” *vs*. “bright”) was significant even within isogenic populations. Representative images of the time course of whole-body GFP expression driven by the *hsp-16*.*2* promoter in the absence of heat shock in three individual animals were shown. Scale bars are 150 µm. (b) The percentage of “bright” worms increases over time within populations of worms grown at 20 °C and 25 °C but increases more rapidly at the higher temperature. (c) *hsp-16*.*2p::*GFP animals show decreased lifespan at 25 °C (30% compared to animals grown at 20 °C, p<0.001, Log-rank test). (d) The fraction of worms that died on each day is temperature-dependent, with animals dying earlier when maintained at 25 °C compared to 20 °C. (e, f) Survival analysis of animals selected on Day 4 to be in the top 1% (brightest 1% of cohort) or top 10% brightest 10% of cohort *vs*. randomly selected control animals form the same cohort. Worms with higher early *hsp-16*.*2p::*GFP expression had significantly shorter remaining lifespans than randomly selected controls (19% shorter for high expression, p<0.001 for the top 1%; 9% shorter for high expression, p<0.05 for the top 10%, Log-rank test). (g) Comparison of detergent-insoluble protein in high and low *hsp-16*.*2p::*GFP expression groups using SDS-PAGE. (h) Detergent-insoluble protein was significantly more abundant in Day 10 animals selected to be in the high expression group (top 10%) than in age-matched, randomly selected control from the same cohort (fold-difference of 1.7±0.1 (mean±SD, p<0.05, paired t-test).

## 2. Experimental Procedures

See also Supporting Information for additional details and data on experimental procedures.

### 2.1. Studies of detergent-insoluble protein at different temperatures

*C. elegans* embryos were hatched on peptone-free NGM plates at 20 °C and transferred to 100 μM 5-fluorodeoxyuridine (FUdR) plates on Day 3 to prevent progeny development. Worms were maintained on 20 °C until Day 8 of life before being transferred to 10, 20, and 25 °C, respectively. Worms were harvested on Day 8 for 20 °C and Day 12 for all temperatures.

### 2.2. Measurement of insoluble protein burden

Our method to quantify the insoluble protein burden in *C. elegans* is based on that reported by David *et al*., with slight modifications (David et al., 2010). Nematodes of about 500 – 3000 individuals were harvested and centrifuged in ice-cold M9 buffer for each condition and harvesting time point. Worm pellets were re-suspended in ice-cold RIPA lysis buffer (50 mM Tris pH 8.0, 150 mM NaCl, 5 mM EDTA, 0.5% SDS, 0.5% SDO, 1% Triton X-100, 1 mM PMSF, Roche cOmplete Mini Protease Inhibitors 1x), and homogenized (Precellys Evolution, Bertin Instruments, France). Total proteins were detected in the gel by silver staining using the Pierce Silver Stain Kit (Life Technologies). Gels were imaged using a Syngene imager (Cambridge, UK). The absolute intensity of proteins detected on silver-stained gels was quantified using ImageJ (NIH, USA). See Supporting Information and fig. S7 for details.

### 2.3. Quantification of intestinal punctae *in vivo*

The number of intestinal polyQ punctae was counted in live on Day 15 using florescent images of live *C. elegans* AM141 [*unc-54p*::Q40::YFP]). Animals were imaged using a microscope camera Leica DFC3000G (Leica M205 with the fluorescence filter set, Leica Microsystems, Germany). Operators were blinded to treatment condition while punctae were counted.

### 2.4. Lifespan measurements

Worms were observed under a light microscope and gently prodded with the worm pick for survival scoring. Individuals that did not respond to gentle prodding were scored as dead. Worms that had crawled off the agar or had suffered from internal hatching were excluded from the analysis. Animals that were not temperature-induced sterile were transferred daily to prevent overcrowding by the progeny. Lifespan analysis was carried out using Online Application for Survival Analysis (OASIS 2, see Table S1 in Supporting Information).

### 2.5. Sorting and measurement of activation of *hsp-16*.*2p*::GFP

For lifespan studies, all worms were sorted manually using a fluorescence microscope (Leica M205 with the GFP fluorescence filter set, Leica Microsystems, Germany). The inclusion criteria being the most significant observable fluorescence of worms in a population defined as “bright” and insignificant observable fluorescence defined as “dim”. While we note that there was a full spectrum of GFP expression ranging from no expression to extremely high expression on the day of sorting, only the brightest or dimmest observed worms in the population were selected for lifespan assays. When sorting worms by the relative degree of GFP induction, the visually brightest individual worms were picked until the required fraction (1% or 10%) of the cohort had been picked. All worms grown at 25 °C were sorted on Day 8 of life, while worms grown at 20 °C were sorted on Day 10 of life. About 100 worms were sorted for high and low expression groups from a total population of 1,000 or 10,000 worms to meet the respective sorting criteria of 10% and 1% of the cohort. For experiments involving total population fluorescence dynamics, any observable fluorescence (regardless of the magnitude of expression) was defined as “bright” while only worm invisible under fluorescence were scored as “dim”. Worms grown on NGM plates were imaged daily on a microscope camera Leica DFC3000G (Leica M205, Leica Microsystems, Germany). For experiments that required worms to be anesthetized, worms were immersed in 1 M levamisole on an agar pad for 3 minutes and directly observed under the microscope. Quantification of worm fluorescence was performed using ImageJ (NIH, USA) and normalized to background fluorescence.

### 2.6. RNAi feeding method

RNAi plasmids were obtained from the Ahringer RNAi library (Source Bioscience), and all clones used were validated by sequencing. The empty vector (EV) L4440 in *E. coli* HT115 (DE3) bacteria was served as a negative control. RNAi bacteria were grown overnight in LB liquid medium containing 12.5 µg/ml tetracycline and 100 µg/ml carbenicillin. NGM containing 100 µg/ml carbenicillin and 1 mM β-D-isothiogalactopyranoside (IPTG) were seeded with RNAi cultures. The RNAi bacteria were harvested using centrifugation (at 4,000 g, 10 minutes at room temperature) and re-suspended to 10^10^ cfu/ml. Embryos were hatched directly on induced RNAi bacteria and grown at 17 °C before being transferred to 25 °C post-L4 stage. For *rsks-1* RNAi treatment, RNAi feeding bacteria was cultured and mixed with EV and *rsks-1* RNAi culture at half a volume each.

### 2.7. Rapamycin treatment

Exposure to rapamycin was carried out, as reported previously (Admasu et al., 2018). Briefly, worms were synchronized using standard hypochlorite treatment and transferred to NGM plates supplemented with 100 µM rapamycin or the vehicle as control at Day 3, and maintained at 25 °C.

### 2.8. MitoSOX Red measurement in live nematodes

Worms were transferred into each well of an opaque 96-well microtiter plate containing 100 μL of M9 buffer. A final concentration of 10 µM MitoSOX in 0.2% DMSO was added to each well. Fluorescence intensity was quantified every 2 minutes for 14 hours using a Hidex Sense plate reader (Hidex, Turku, Finland) at room temperature using an excitation wavelength of 396 nm and an emission wavelength of 579 nm.

### 2.9. Statistical analysis

All data were generated and analyzed in GraphPad Prism (v 5.02). Data are presented as means ± error bar (either SEM or SD) unless noted otherwise. Values of p<0.05 (*) were considered statistically significant.

## 3. Results

### 3.1. Age-dependent changes in insoluble protein level

Insoluble protein, specifically below 45 kDa molecular weight, accumulates with age in *C. elegans* (David et al., 2010). We established and validated a variant of this assay in our laboratory (see Experimental Procedures and Supporting Information). To determine if our modified method was able to detect the previously reported age-dependent increase reliably, we first compared the relative abundance of low-molecular-weight (below 49 kDa) insoluble protein between young (Day 4) and aged (Day 12) animals (Fig. S1**a**). To generate age-synchronized worm cohorts for this study, we utilized the germline-deficient strain, JK1107 (*glp-1(q224) III*). At the restrictive temperature of 25 °C, JK1107 animals are defective in germline proliferation and do not lay eggs (Austin & Kimble, 1987). This renders them sterile, avoids overgrowth with progeny, and prevents inadvertent addition of progeny-derived material to samples taken from the aging cohort. We collected replicate samples from independent cohorts, sampled on Day 4 and Day 12. For each sample, the abundance of insoluble protein, relative to total soluble protein, was quantified, and results were averaged over three independent biological repeats for each age group (Fig. S1**b**). Statistical analysis of the relative amount of insoluble protein, normalized to soluble protein, confirmed the initial visual impression that insoluble protein increased significantly relative to soluble protein between Day 4 and Day 12 (fold change of 2.5±0.5 (mean±SD), p<0.05, paired t-test). These data confirmed that age-dependent changes can be reliably detected using this assay based on only three biological repeats (Fig. S1**b**) and are consistent with previous observations showing that insoluble proteins increase with age (David et al., 2010; Groh et al., 2017; Reis-Rodrigues et al., 2012).

Oxidative damage to protein can drive protein aggregation (Reeg & Grune, 2015). We therefore asked if the insoluble protein fraction was, on average, more oxidized than soluble protein. Protein carbonyl content (PCC), a form of oxidative protein damage, has been shown to increase with age in *C. elegans* and is commonly used as endpoint assay of oxidative protein damage (Adachi, Fujiwara, & Ishii, 1998; Goto et al., 1999; Gruber et al., 2011). Consistent with previous observations, we confirmed a significant increase in PCC between Day 4 and Day 12 of age in soluble protein (1.8±0.4 fold change (mean±SD), p<0.05, t-test, fig. S1**c**). To test whether there was a significant difference between soluble and insoluble protein in terms of the degree of oxidative damage, we normalized PCC values for each experiment to the average PCC of soluble protein at that age and then pooled the normalized PCC for Day 4 and Day 12 (Figure S1**c**). Comparing the relative degrees of oxidative damage in the soluble and insoluble fractions revealed that, on average, insoluble protein was significantly more oxidized than soluble protein (p<0.01 for both comparisons, paired t-test, fig. S1**d**).

Interestingly, while oxidative damage overall and in the soluble fraction increases with age, there was no such age-dependent increase in the degree of protein oxidation within the insoluble fraction between Day 4 and Day 12 (Fig. S1**c**). The age-dependent global increase in protein oxidation, therefore, seems to be driven by increased oxidation of protein in the soluble fraction and by the increasing amount (but not the degree of oxidation) of insoluble protein.

### 3.2. Temperature modulates lifespan and insoluble protein level

*C. elegans* are ectotherms, meaning that their body temperature and metabolism depend on environmental temperature. Body temperature impacts the rate of many metabolic processes, including protein turnover and stability (Becktel & Schellman, 1987). In ectotherms, including *C. elegans* (Klass, 1977) and *Drosophila melanogaster* (Economos & Lints, 1986), temperature and lifespan are typically inversely related. However, while exposure to stressfully high temperatures (heat shock) rapidly causes an increase in unfolded protein and triggers adaptive responses (Nakai, 2016), the effects of moderate, chronic changes in temperature on insoluble protein dynamics are more complex. Protein synthesis and degradation (turnover) rates increase with temperature, but so do rates of protein unfolding and the production rates of damaged protein, typically as side-products of metabolism (Schulte, 2015).

We first evaluated the lifespan of WT nematodes cohorts, maintained throughout their entire lifespan at 10, 14, 16, 20, and 25 °C (Fig. S2**a**). In agreement with previous publications on *C. elegans* lifespan (B. Kim, Lee, Kim, & Lee, 2020; Klass, 1977), we found a strong influence of temperature on lifespan, with cohorts grown at lower temperatures living significantly longer, while worms that were grown at higher temperatures living shorter (Fig. S2**a**). We next determined the rate of accumulation of insoluble protein in nematodes maintained at different temperatures. For insoluble protein assays, animals were maintained until mid-adulthood (Day 8) at 20 °C, after which they were transferred to different temperatures. This was done to avoid potential confounding effects of temperature on their developmental schedule. Under these conditions, animals spent a smaller fraction of their life at different temperatures and lifespan differences were less pronounced than for animals maintained at different temperatures throughout life (Fig. S2**b**). However, animals maintained at a higher temperature were still short-lived. We then assessed changes in insoluble protein between Day 8 and Day 12 of life as a function of temperatures (Fig. 1**b**, fig. S2**c** for full gel). We normalized the detergent-insoluble protein on Day 12 for each temperature (10, 20, and 25 °C) to the average detergent-insoluble protein level on Day 8 of control (20 °C). In animals consistently maintained at 20 °C, we observed only an insignificant trend towards a further age-dependent increase in insoluble protein over the four-day period between Day 8 and Day 12 (fold change of 1.4±0.3 (mean±SD), p=0.16, t-test, normalized to Day 8). On Day 12, the means of relative detergent-insoluble protein in each of the temperatures tested were significantly different (p<0.05, one-way ANOVA). Post-tests showed that animals shifted to 25 °C experienced a significantly more rapid increase in insoluble protein between Day 8 and Day 12 (fold change of 2.2±0.3 (mean±SD), p<0.05 for both 25 °C versus 10 °C and 25 °C versus 20 °C, one-way ANOVA, fig. 1**b**). Due to the need to initially grow animals at non-restrictive temperatures, we did not use JK1107 for this study. Instead, WT animals were grown on plates supplemented with FUdR to prevent contamination of progeny.

We next utilized a *C. elegans* strain (AM141) expressing a YFP labeled form of a specific aggregation-prone protein. This strain has been developed to study protein aggregation driven by polyglutamine (polyQ) repeats, the underlying cause of Huntington Disease (HD) (Morley et al., 2002). AM141 expresses yellow fluorescent protein (YFP) fused to aggregation-prone glutamine repeats in body wall muscle, enabling visual scoring of polyQ-driven aggregates by counting the number of YFP-fluorescent puncta (Morley et al., 2002). We used the AM141 polyQ strain to directly visualize the dependence of visible aggregates formation on the temperature *in vivo*. Consistent with our more general observations regarding increased global protein aggregation at higher temperatures and with a previous report (Moronetti Mazzeo, Dersh, Boccitto, Kalb, & Lamitina, 2012), we observed a temperature-dependent increase in the formation rate of visible polyQ aggregates in the AM141 strain (Fig. 1**c**). In summary, higher temperatures lead to a shorter lifespan and cause more rapid accumulation of insoluble and aggregated protein, and this was true for both endogenous *C. elegans* protein and specific, disease-related polypeptides.

### 3.3. The impact of the germline on insoluble protein dynamics and lifespan

CB4121 *(sqt-3(e2117) V)* is a *C. elegans* strain carrying a mutation preventing collagen synthesis at high temperatures (Wang, Oakley, Carr, Sowa, & Ruvkun, 2014). Unlike the germline defective *glp-1* strain used for most of our studies, adult *sqt-3* mutants are capable of normal reproduction. Due to the defect in collagen synthesis, cuticle development in larvae is defective, preventing offspring from developing into adults when grown at the restrictive temperature. To test the dependence of insoluble protein on the presence of a functional germline, we compared lifespan and insoluble protein dynamics between age-matched cohorts of the germline-deficient *glp-1* and the germline-competent *sqt-3* strains. The mean lifespan of *sqt-3* mutants was approximately 12% shorter compared to *glp-1* (p<0.001, Log-rank test, fig. 1**d**, Table S1 for all lifespan data statistics). We next examined the accumulation rate of insoluble protein in *sqt-3* mutants and found that, compared to the *glp-1*, insoluble protein accumulated much more rapidly in *sqt-3* between Day 4 and Day 12 (fold increase of 7.7±1.6 (mean±SD) in *sqt-3 vs*. fold increase of 1.4±0.05 (mean±SD) in *glp-1*, p<0.01, 99% CI comparison, t-test, fig. 1**e, f**, fig. S2**d** for full gel). The observation that accumulation of insoluble protein occurs more rapidly in the strain carrying a functional germline is consistent with its shorter lifespan and with previous observations that protein aggregation is slowed in a gonad-less (*gon-2*) background (David et al., 2010).

### 3.4. Higher *hsp-16*.*2* expression in early life predicts shorter remaining lifespan and scales with temperature

All data presented thus far was generated using synchronized cohorts of aging animals, meaning that they are correlative in nature and pertain to population averages. However, even cohorts of nematodes comprising isogenic clones, feeding off the same source of bacterial food and experiencing near-identical environments exhibit significant heterogeneity in individual lifespan and their aging-trajectories (Herndon et al., 2002). We therefore asked if individual aging trajectories and remaining lifespans of individual animals were related to proteostatic failure in these animals. To explore this question, we utilized a transgenic *C. elegans* transcriptional reporter strain (CL2070), which carries a green fluorescent protein (GFP) gene under the control of the *hsp-16*.*2* heat shock promoter (Link et al., 1999). However, even in worms maintained consistently at a non-stressful temperature of 20 °C and never exposed to any heat shock, *hsp-16*.*2* is sporadically activated and such activation becomes more prevalent with age (Fig. 2**a**). As *hsp-16*.*2* is a downstream target of heat shock factor-1 (HSF-1) and DAF-16 and is directly involved in heat shock responses (Haslbeck & Vierling, 2015; Rodriguez, Snoek, De Bono, & Kammenga, 2013), we hypothesized such spontaneous activation of *hsp-16*.*2* might indicate an endogenous crisis in maintaining proteostasis and that this might be predictive of a shorter remaining lifespan.

To test this hypothesis, we first carried out a series of longitudinal and cohort studies, visually scoring spontaneous *hsp-16*.*2* activation in CL2070 animals as a function of age. On each day, we compared young reference animals showing no or minimal GFP expression with animals from aging cohorts, scoring only those animals as “bright” that showed clearly detectable GFP signal. Scoring was done by toggling bright-field illumination on and off and scoring presence or absence of fluorescence. We used WT animals carrying no GFP transgene as negative control and young, and heat-shocked CL2070 animals with robust induction of the reporter as positive control. Animals from the aging cohort were scored as “dim” when they were invisible under fluorescence and as “bright” if fluorescence was clearly present. Data from this study confirm that old animals are indeed more likely to show spontaneous *hsp-16*.*2* activation than young animals (Fig. 2**a, b**).

This study revealed that almost all animals show some degree of spurious/aberrant activation at some point prior to death (Fig. 2**b**). As expected, animals that were grown at 25 °C were 30% shorter-lived, and these animals also tended to spontaneously express *hsp-16*.*2* earlier than those maintained at 20 °C (Fig. 2**b-d)**. The magnitudes of change in GFP dynamics were comparable with the change in mean lifespan. At 25 °C, lifespan was reduced by six days, while the day where 50% of the population showed significant *hsp-16*.*2* activation occurred four days earlier than at 20 °C.

To directly test if this spontaneous *hsp-16*.*2* activation was associated with lifespan detriments, we next identified young animals that showed unusually early and robust activation of *hsp-16*.*2*. This was done by visually sorting animals on Day 8 of life, selecting animals that showed clear GFP fluorescence (see Experimental Procedures). Using large cohorts, we selected the visually brightest 1% or 10% of animals and transferred these to separate lifespan cohorts. Control sample cohorts showed no increased GFP expression and were picked randomly (under bright-field conditions where GFP is invisible) from the same source plates. Using this approach, we found that animals showing higher *hsp-16*.*2* expression on Day 8 of age had a significantly shorter remaining lifespan than controls randomly selected from the same cohort. The brightest 1% of the cohort had a lifespan that was, on average, 19% (mean lifespan) shorter than the cohort average (p<0.001, Log-rank test, fig. 2**e**). Even the brightest 10% of the cohort still suffered a statistically significant 9% reduction in mean lifespan (p<0.05, Log-rank test, fig. 2**f**).

Previous evidence shows that transgenic overexpression of *hsp-16*.*2* itself, or its induction in response to hormetic heat shock results in lifespan *extension* rather than shortening (Cypser & Johnson, 2002). This suggests that it is not an expression of *hsp-16*.*2* or its GFP reporter construct itself that is detrimental to CL2070 survival, but that *hsp-16*.*2* activation is indicative of an intrinsic crisis state associated with shorter remaining lifespan. These observations are therefore consistent with the hypothesis that spontaneous *hsp-16*.*2* transcriptional activation is indicative of an endogenous crisis state. Importantly, we never observed a reversal of this phenotype (recovery), with animals inevitably dying within 3-8 days of showing spontaneous activation (Fig. S3).

Next, we asked if spontaneous activation of *hsp-16*.*2* was indeed indicative of increased insoluble protein burden and failure in proteostasis. To test this hypothesis, we performed insoluble protein measurements on samples taken from either the “bright” or “dim” subsets of the aging worm cohort. Using the same sorting procedure outlined above (using 10% as the cutoff in this case), and comparing them to randomly-selected controls from the same cohort, we were able to confirm that animals showing early *hsp-16*.*2* activation exhibited significantly higher accumulation of insoluble protein than animals without such activation (fold difference of 1.7±0.1 (mean±SD), p<0.05, paired t-test, fig. 2**g, h**). This observation suggests that “bright” worms activate heat shock pathways early and experience higher levels of protein damage or aggregation than age-matched “dim” animals. Because the same animals are also short-lived, this observation supports a link between the stochastic failure of proteostasis, protein aggregation, and shorter lifespan in individual nematodes (Fig. S3).

### 3.5. Treatment with mTOR inhibitor, rapamycin, delays *hsp-16*.*2* expression and reduces insoluble protein

The mTOR pathway is known to control key processes underlying protein synthesis and proteostasis, and the eponymous mTOR inhibitor, rapamycin, extends lifespan in WT *C. elegans* at 20 °C (Robida-Stubbs et al., 2012). We performed lifespan analysis on CL2070 *hsp-6*.*2p*::GFP animals grown at 25 °C with and without rapamycin treatment and found that the lifespan of these animals was also extended by rapamycin (p<0.05, Log-rank test, fig. 3**a**). While rapamycin significantly extended the lifespan of CL2070 animals, maintained at 25 °C throughout life, the resulting lifespan increase was only 9%, which is smaller than benefits that are typically seen in WT animals maintained at 20 °C (Admasu et al., 2018; Robida-Stubbs et al., 2012). We then compared levels of insoluble protein between populations treated with rapamycin and control. We found the insoluble protein levels were borderline lower in populations treated with rapamycin (fold change of 0.5±0.03 (mean±SD), p=0.06, t-test) suggesting that mTOR inhibition by rapamycin was able to delay age-dependent insoluble protein (Fig. 3**b, c**, fig. S4 for full gel).

**Figure 3.**
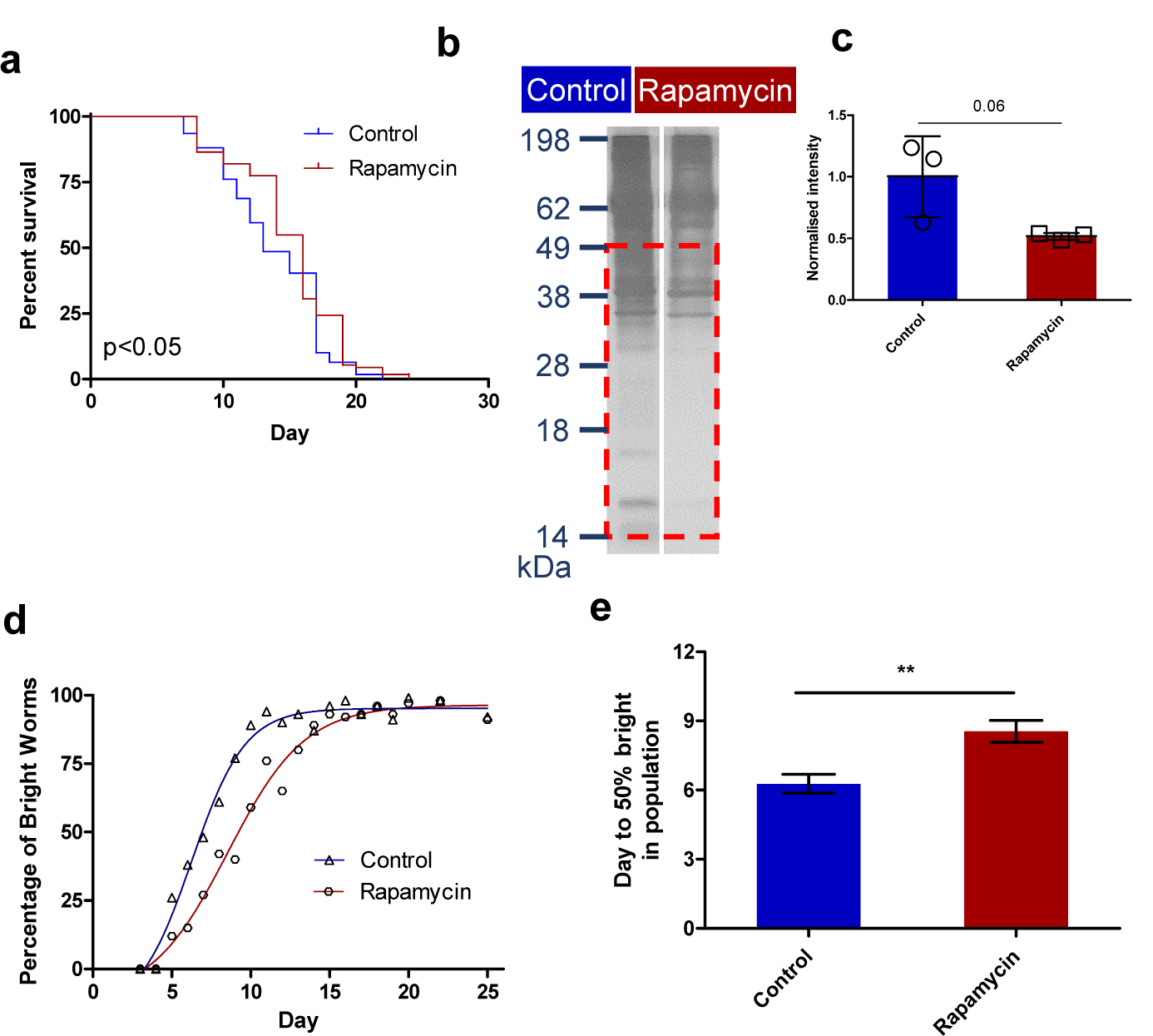
Rapamycin delays the spurious mid-adulthood expression of *hsp-16*.*2p*::GFP and reduces the levels of detergent-insoluble protein. (a) Survival curves of *hsp-16*.*2p::*GFP worms grown at 25 °C with or without (control) rapamycin. Rapamycin (100 µM) extended the lifespan of *hsp-16*.*2p::*GFP nematodes at 25 °C compared to control (9%, p<0.05, Log-rank test). (b) Representative gel image of insoluble protein from Day 8 worms with and without exposure to rapamycin. (c) Quantification of insoluble protein levels of rapamycin-treated worms normalized to the average of soluble protein and control of each condition showed a trend towards lower levels of insoluble protein in treated animals with a fold difference of 0.5±0.03 (mean±SD) that did, however, not reach statistical significance (p=0.06, t-test). (d) The proportion of “bright” worms over time within a population of worms with or without rapamycin. (e) Analysis of non-linear curves fitting showed that the difference between the time course of age-dependent *hsp-16*.*2p::*GFP for control (6.3±0.4, df=16) and rapamycin-treated animals (8.6±0.5, df=16) is statistically significantly different (p<0.01, t-test, Table S2 for statistical analysis).

To confirm that pharmacological inhibition of the mTOR pathway delayed the age-dependent proteostatic crisis, we next tested if rapamycin was able to alter the time course of spontaneous *hsp-16*.*2* activation. Using the same approach as above, we found that exposure to rapamycin indeed delayed *hsp-16*.*2* induction in CL2070 (Fig. 3**d**). When time courses for *hsp-16*.*2* activation were compared for treated and untreated cohorts, 50% of the untreated animals showed visible *hsp-16*.*2* activation on Day 6. In comparison, rapamycin-treated animals reached this level of induction only after Day 8 of life (average days to 50% “bright” in populations: 6.3 (control) *vs*. 8.6 (rapamycin), p<0.01, t-test, fig. 3**e**, Table S2). These observations indicate that the mTOR inhibition by rapamycin benefited the *hsp-16*.*2p*::GFP worms resulting in reduced accumulation of insoluble protein, delayed the late-life crisis in proteostasis, as measured by activation of the *hsp-16*.*2* promoter and extended lifespan.

### 3.6. Inhibition of ribosomal S6 kinase reduces insoluble protein

A key downstream target of rapamycin is the ribosomal kinase S6K, with mTOR inhibition reducing protein translation through S6K inhibition. Recent evidence suggests that reduced protein translation has age-modifying and lifespan-extending effects, likely due to improved proteostasis (King et al., 2008; Zhao et al., 2015), even when initiated late in life (Solis et al., 2018). We therefore asked if insoluble protein was directly affected by RNAi against S6K/*rsks-1*. Given the effect of the germline on insoluble protein (Fig. 1**f, g**) and lifespan (David et al., 2010; Hsin & Kenyon, 1999), we first confirmed *rsks-1* was sufficiently inhibited by RNAi feeding using qPCR in *glp-1* mutants and N2 (Fig. S5**a**), and then lifespan that *rsks-1* RNAi results in lifespan extension in WT as well as in *glp-1* and *sqt-3* mutants at 25 °C. Two of these strains have an intact germline (WT and *sqt-3*), while *glp-1* does not (Austin & Kimble, 1987; Wang et al., 2014). Interestingly, *rsks-1* RNAi significantly extended the mean lifespan of both germline-competent strains, extending mean lifespan of WT by 5% (p<0.05, Log-rank test, fig. 4**a**) and by 9% in *sqt-3* mutants (p<0.01, Log-rank test, fig. 4**b**). However, in the *glp-1* background, lifespan was not statistically significantly affected in our hands (p=n.s., Log-rank test, fig. 4**c**). These data suggest that both germline-competent strains (WT and *sqt-3)* benefit more from reduced translation than the germline-deficient *glp-1*. Because WT animals produce a large number of live offspring, interfering with the insoluble protein assay, we used *sqt-3* instead of WT as the germline-competent strain for our subsequent insoluble protein studies. We investigated insoluble protein with and without RNAi-mediated *rsks-1* knockdown in aging *sqt-3* and *glp-1* mutants (Fig. 4**d, e**, fig. S5**b, c** for full gel). Surprisingly, despite its lack of benefit on lifespan, the rate of accumulation of insoluble protein was significantly reduced in *glp-1* mutants fed with *rsks-1* RNAi compared to control (fold difference of 0.7±0.02, p<0.01, paired t-test, fig. 4**e**), while it remained unchanged in *sqt-3* mutants (fold difference of 1.2±0.03, p=n.s, paired t-test, fig. 4**e**).

**Figure 4.**
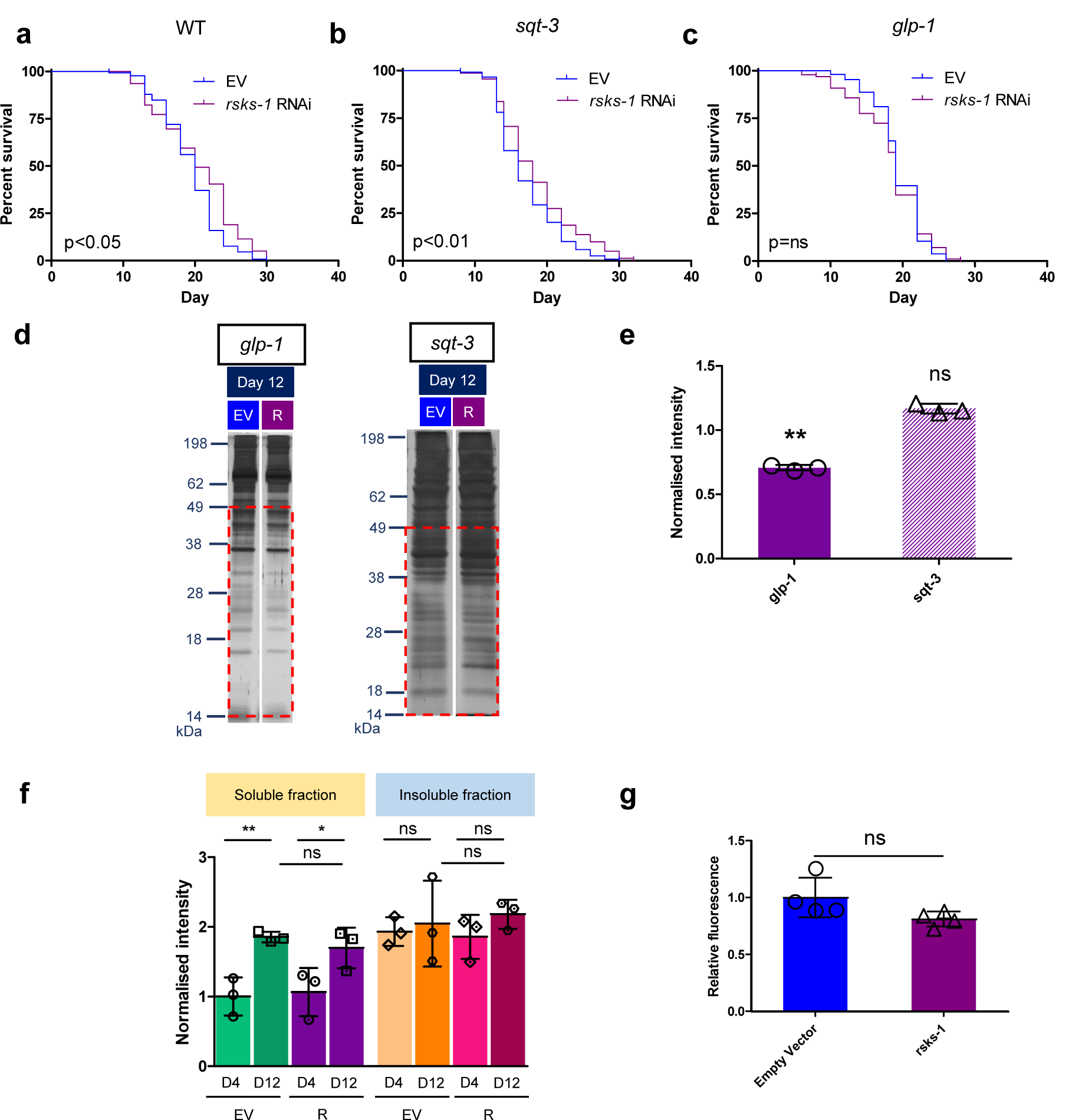
Reduction of translation by downregulating *rsks-1* slower the accumulation of insoluble protein. (a-c) Survival curves of (a) WT and (b) *sqt-3* and (c) *glp-1* mutants treated with *rsks-1* RNAi. Reduction of *rsks-1* significantly increased lifespan in WT (p<0.05) and *sqt-3* (p<0.01) but not in *glp-1* (p=n.s, Log-rank test). (d) Representative gel images of *rsks-1* RNAi-treated *glp-1* and *sqt-3* populations on Day 12. (e) Measurement of the detergent-insoluble protein in *rsks-1* RNAi animals of Day 12 normalized to control (empty vector, EV) showed a lower burden of insoluble protein in *glp-1* animals (fold difference of 0.7±0.02 (mean±SD), p<0.01, paired t-test) but unchanged in *sqt-3* (fold difference of 1.2±0.03 (mean±SD), p=n.s, paired t-test). (f) Comparison of normalized PCC measurement of SF and IF compared between Day 4 and Day 12 for control and *rsks-1* RNAi-treated *glp-1* animals. Densitometry measurements of all groups, including SF and IF, were normalized against the mean of SF of Day 4 EV. When comparing PCC between Day 4 and Day 12 of age, there was a comparable increase in the SF of both EV (fold change of 1.9±0.08 (mean±SD), p<0.01, t-test) and *rsks-1* knockdown animals (fold change of 1.7±0.3 (mean±SD), p<0.05, t-test). For the IF, the results from densitometry were normalized against the mean of SF of Day 4 of EV. One-way ANOVA analysis showed that there was no significant difference between IFs of Day 4 and Day 12. (g) There was only a slight, non-significant trend towards a decrease in MitoSOX fluorescence in cohorts of *rsks-1* RNAi-treated animals relative to EV on Day 8 of age (fold difference of 0.8±0.07 (mean±SD), p=0.09, t-test). Data represented as four biological replicates comprising 100 individual nematodes in each bar.

### 3.7. Reduction of *rsks-1* does not reduce oxidatively damaged protein or endogenous oxidative damage

At 25 °C, germline-deficient *glp-1* mutants live longer and accumulate insoluble protein more slowly than strains with functional germline (Fig. 1**e, f**), consistent with a detrimental role of insoluble protein on lifespan. However, *rsks-1* RNAi does not further extend the lifespan of *glp-1* mutant animals, despite further reducing accumulation of insoluble protein (Fig. 4**a-e**). Given our evidence that insoluble protein is more oxidized than soluble protein, we wondered if changes in the relative abundance of oxidatively damaged protein might explain this apparent contradiction. We therefore compared the age-dependent accumulation of oxidized protein in the soluble and insoluble fraction of *glp-1* mutants with and without *rsks-1* RNAi feeding. As before, we observed a significant age-dependent increase of PCC in soluble protein for both EV (fold change of 1.9±0.08 (mean±SD), p<0.01, t-test) and *rsks-1* RNAi (fold change of 1.7±0.3 (mean±SD), p<0.05, t-test). However, there was no significant difference between EV and *rsks-1* RNAi-treated animals (Fig 4**f** and fig. S5**d** for full blot).

Consistent with this lack of change in oxidized protein, there was no significant difference in reactive oxygen species (ROS) production between controls and animals in animals fed on *rsks-1* RNAi (fold difference of 0.8±0.07 (mean±SD), p=0.09), as assessed using the MitoSOX Red fluorescence (Fig. 4**g**). Together, these data suggest that the reduction of insoluble protein in the *rsks-1* RNAi-treated animals was not due to reduced ROS production and did not result in a lower burden of oxidatively damaged protein.

## 4. Discussion

Damage accumulation theories suggest that aging is a consequence of the accumulation of macromolecular damage, with over 300 different forms of damage implicated in aging (Viña, Borrás, & Miquel, 2007). However, amongst these types of damage, accumulation of unfolded, terminally modified, insoluble and aggregated protein is considered a strong candidate for a “public”, that is, evolutionarily highly conserved mechanisms of aging (David et al., 2010; Partridge & Gems, 2002).

Consistent with previous reports, we found evidence for a clear age-dependent accumulation of insoluble protein under different experimental conditions in all of the strains that we tested (David et al., 2010; Groh et al., 2017; Reis-Rodrigues et al., 2012). Insoluble protein was, on average, more oxidized than soluble protein. However, while there was also an age-dependent increase in oxidative damage in detergent-soluble protein, as reported previously (Adachi et al., 1998; Brys, Vanfleteren, & Braeckman, 2007; Goto et al., 1999; Gruber et al., 2011), there was no further age-dependent increase in the degree of oxidative damage of protein in the insoluble fraction. Our data is consistent with the notion that ROS-mediated damage is one driver of protein aggregation and suggests an age-dependent collapse in proteostasis that coincides with disruption in the balance between damage generation and damage removal by turnover (Hipp et al., 2019; Levy et al., 2019; Weids & Grant, 2014).

Loss of germline is known to extend lifespan in *C. elegans* (Hsin & Kenyon, 1999; Kenyon, 2010), and consistent with this, it has been reported that accumulation of insoluble protein is slower in the absence of germline (David et al., 2010). In agreement with these reports, we find that the germline-deficient strain had a longer lifespan than either of the two strains with functional germline (Fig. 1**e**) and that it accumulated insoluble protein more slowly (Fig. 1f, **g**). These observations confirm the notion that germline signaling negatively affects proteostasis, possibly due to the fact that maximizing offspring requires consistently rapid protein synthesis (Khodakarami et al., 2015) and suggests that this process is detrimental for lifespan. Furthermore, FUdR has been reported to improve proteostasis (Feldman, Kosolapov, & Ben-Zvi, 2014) and may block some of the lifespan extension in *C. elegans* maintained at 10 and 15 °C (Lee et al., 2019). This confounding effect of FUdR might explain why we did not observe more pronounced differences in insoluble protein between 10 and 20 °C in FUdR treated cohorts (Fig. 1**b**).

We further found that spontaneous activation of transcription from the *hsp-16*.*2* promoter increases with age and that such activation signals a proteostatic crisis associated with a higher burden of insoluble protein and shorter remaining lifespan in individual animals. The fact that we never observed crises resolution or recovery once this spontaneous activation of the *hsp-16*.*2* promoter had occurred (Fig. S3), suggests that proteostatic failure signals an endogenous and irreversible physiological crisis in aging nematodes. This crisis may be similar to some age-dependent diseases in humans, driven by aggregation of endogenous proteins, occur stochastically late in life, and generally do not spontaneously resolve.

We found that treatment with rapamycin delayed the spontaneous activation of the *hsp-16*.*2* promoter but found only a trend towards reduced age-dependent insoluble protein, at least in *glp-1* mutants. This may, in part, be explained by the fact that *glp-1* mutants, lacking a germline, already have lower insoluble protein levels and are comparatively long-lived (David et al., 2010). However, based on prior observations that inhibiting translation extends lifespan and improves proteostasis in *C. elegans* (Solis et al., 2018), we next silenced the *C. elegans* ribosomal kinase S6K using RNAi and were able to demonstrate a significant reduction in cohort-level insoluble protein in both germline-deficient *glp-1* and germline-competent *sqt-3*. Interestingly, this intervention, again, had limited lifespan benefits in *glp-1* but had a more significant benefit in the presence of a function germline (both in *sqt-3* and WT). These data indicate that in *C. elegans* with functional germlines, proteostasis fails earlier and more severely and that under such conditions, reduced translation is especially beneficial. This may be a generalized example of a previous observation that overproduction of yolk proteins and embryonic growth drive accumulation of abnormal protein and age-dependent decline (Ezcurra et al., 2018). This intervention did not affect levels of oxidatively damaged protein, consistent with the idea that hyperfunction directly contributes to age-dependent failure and disease (Blagosklonny, 2012; Ezcurra et al., 2018; Gems & de la Guardia, 2013). An important observation in this context is the surprisingly large role of intrinsic randomness in the dynamics of proteostatic failure in individual nematodes. Preceding the large stochastic variability in individual lifespan, we observe that, even in synchronized cohorts of isogenic *C. elegans* maintained under standardized conditions, the critical time point at which proteostatic crisis first occurs is highly random, illustrating that important aspects of individual aging trajectories are driven by intrinsic biological stochasticity. This suggests that random events that occur well before the onset of any overt age-dependent dysfunction control the late-life fate of individual animals. In other words, aging is not just a product of interactions between genes and the environment, but random events at the cellular level contribute significantly to individual aging trajectories.

## Declarations

### Funding

This study was supported by the Ministry of Education Singapore and Yale-NUS College (R-607-000-348-114, R-607-265-355-121 and R-607-000-383-114), and NUS Yong Loo Lin School of Medicine Research Scholarship.

### Conflicts of interest

We declare that there is no conflict of interest.

### Ethic approval/Consent to participate/Consent for publication

Not applicable.

### Availability of data and material

The data that supports the findings of this study are available in the supplementary material of this article.

### Code availability

Not applicable.

### Author Contributions

YZ, SHYL, and JG designed the study and wrote the manuscript. YZ and SHYL performed most experiments and analyzed results. NLF and JG participated in the interpretation of the data. All authors commented and approved the manuscript.

## Acknowledgments

All *C. elegans* strains reported in this article were provided by the Caenorhabditis Genetics Centre, which is funded by the NIH Office of Research Infrastructure Programs (P40 OD010440). We thank Dr. Scott Leiser for kindly provided the empty vector RNAi control clone.

## Notes

### Competing Interest Statement

The authors have declared no competing interest.

