## Supplementary File for "Inhibition of mTOR decreases insoluble protein burden by reducing translation in *C. elegans*"

**SUPPORTING INFORMATION**

**Experimental Procedures**

**Nematode strains and maintenance**

The *C. elegans* strains, WT N2 (Bristol), AM141 (*(rmIs133)* [*unc-54p*::Q40::YFP]), CB4121 (*sqt-3(e2117)*), CL2070 (*(dvIs70*) [*hsp-16.2p::*GFP; *rol-6(su1006)*]), and JK1107 (*glp-1(q224)*), were maintained using standard methods (Stiernagle, 2006) for the experiments. The preparation of nematode growth media (NGM) was with the addition of streptomycin (Sigma) at the final concentration of 200 µg/ml. Streptomycin-resistant bacteria strain (*Escherichia coli* OP50-1), unless specified, was added as a food source to the solid media at 10^10^ cfu/ml. Nematodes were cultured at 20 °C unless otherwise stated, and they were grown to the gravid adult stage before synchronization of age using the standard bleaching treatment. JK1107 and CB4121 animals were maintained at the restrictive temperature (25 °C) to prevent progeny.

| Strains used in this Study | | |
| --- | --- | --- |
| **Name** | **Genotype** | **Strain Origin** |
| N2 | Wild-type (WT) | CGC |
| CL2070 | *dvIs70*[*hsp-16.2p::*GFP; *rol-6(su1006)]* | CGC |
| JK1107 | *glp-1(q224) III* | CGC |
| CB4121 | *sqt-3(e2117) V* | CGC |
| AM141 | *unc-54p*::Q40::YFP | CGC |

**UV inactivation of *E. coli* OP50-1**

For temperature studies, *E. coli* bacteria culture was then pipetted onto a peptone-free NGM plate, and the seeded plates were exposed to ultraviolet light at 1000 mJ/cm^2^ using a UV crosslinker, SpectroLine XLE-1000/FB (Spectronics Corp., Westbury, NY).

**Sequential protein extraction in RIPA lysis buffer and urea lysis buffer**

Detergent-soluble and -insoluble protein extraction methods were adapted from (David *et al*., 2010). Each tube consisted of worms and lysis buffer of 400 μl, and ceramic beads were added subsequently, and the worms were homogenized using a bead-beater tissue homogenizer. Every homogenization cycle was to homogenize (H) for 20 seconds and to pause (P) for 10 seconds, and this cycle was repeated for three times for a complete extraction sequence (H: 20s, P: 10s, x3). The lysates were centrifuged to collect insoluble pellets at 14,000 g for 10 minutes, at 4 °C, and 300 μl of the detergent-soluble protein extract was transferred into a fresh tube and labeled as soluble fraction, leaving the pellet undisturbed with 100 μl of the remaining lysate. The pellet was re-extracted with the addition of 300 μl of fresh RIPA lysis buffer. This lysate was homogenized with the same sequence mentioned (H: 20s, P: 10s, x3) and centrifuged for another five times to remove detergent-soluble protein (labeled as soluble wash 1-5). Subsequently, 300 μl of urea lysis buffer (8 M Urea, 2% SDS, 1 mM DTT, 50 mM Tris pH 7.4) was added the pellet with 100 μl of RIPA lysis buffer at room temperature to extract detergent-insoluble protein (homogenization sequence, H: 20s, P: 10s, x2). The lysates were centrifuged at 14,000 g for 10 minutes at room temperature. The first fraction of detergent-insoluble protein, 300 μl of the lysate was transferred to a new tube and labeled as insoluble fraction. Another 300 μl of fresh urea lysis buffer was added to the pellet and the remaining 100 μl lysate. The homogenization and centrifugation steps were repeated for another five cycles (labeled as insoluble wash 1-5) to arrive at an insoluble pellet, which was labeled as pellet (Fig. S6).

Lysates from soluble fraction, wash 1-3, were combined to obtain a pooled soluble protein, and a total protein of 2 μg for pooled soluble was required for loading. Pooled soluble protein was loaded to check for equal loading of proteins. Lysates from insoluble fraction, wash 1-3 were combined to get a pooled insoluble protein, and the volume required to load for pooled soluble was multiplied by 15x to obtain the final volume required to load pooled insoluble protein. All samples were loaded with SDS reducing buffer loading dye. Samples were subjected to 12% SDS-PAGE gel, and the proteins were separated in Tris-Glycine SDS running buffer according to Bio-Rad protocol.

RNA extraction and quantitative PCR

RNA was extracted using the QIAGEN RNeasy Mini Kit (QIAGEN, Netherlands), subjected to an additional DNase step using QIAGEN DNase I Kit according to the manufacturer’s protocol. Isolated RNA was proceeded with reverse-transcription PCR using Promega GoScript Reverse Transcriptase (Promega, Wisconsin, USA) following the manufacturer’s instructions to synthesis cDNA templates. qPCR was performed using Applied Biosystems PowerUp SYBR Green Master Mix (Thermofisher Scientific, USA) with the addition of primers (Integrated DNA Technologies, USA) in StepOnePlus Real-Time PCR System (Thermofisher Scientific, USA) for amplification (UDG activation at 50 °C, 2 minutes, polymerase activation at 95 °C, 2 minutes, 40 cycles of denaturing at 95 °C, 3 seconds, annealing/extension at 60 °C, 30 seconds). Gene expression of target genes such as *rsks-1* was analyzed. mRNA levels of target genes were normalized to the mean of the gene *act-1*. Primer sequences are listed in Supporting Table S3.

PCC measurement

For each sample, 2 μg of protein from total soluble fraction and insoluble fraction (15x the volume of the soluble fraction) were derivatized according to manufacturer’s protocol (OxyBlot Protein Oxidation Detection Kit, Merck Millipore, Germany). The derivatized proteins was loaded to a Bio-Dot SF Microfiltration apparatus and transferred to a nitrocellulose membrane (Bio-Rad Laboratories, USA). The membranes were blocked, then probed with an anti-DNPH primary antibody, followed by an anti-rabbit HRP conjugated IgG antibody. Antibody-bound proteins were detected by chemiluminescence using SuperSignal West Dura Extended Duration Substrate (Thermofisher Scientific, USA). Images were taken using a Syngene system and analyzed using ImageJ (NIH, USA).

Day 4

Day 12

Day 4

Day 12

198

Detergent-insoluble

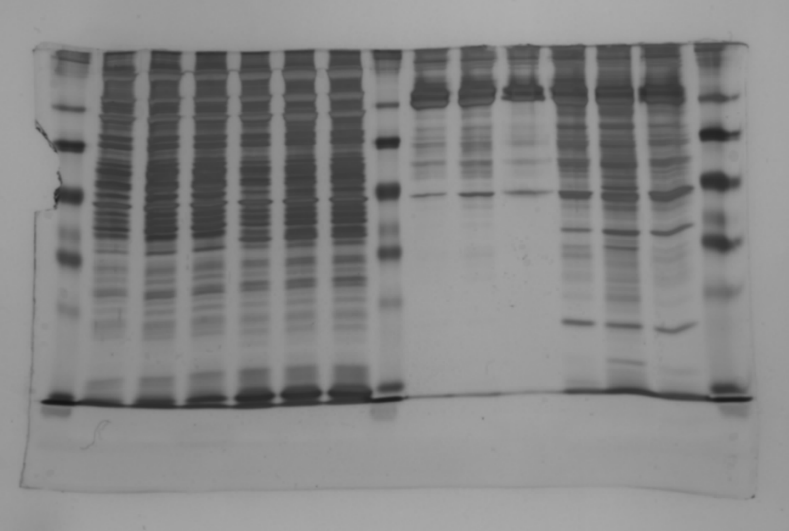

Detergent-soluble

62

48

38

28

18

14

kDa

198

62

48

38

28

18

14

kDa

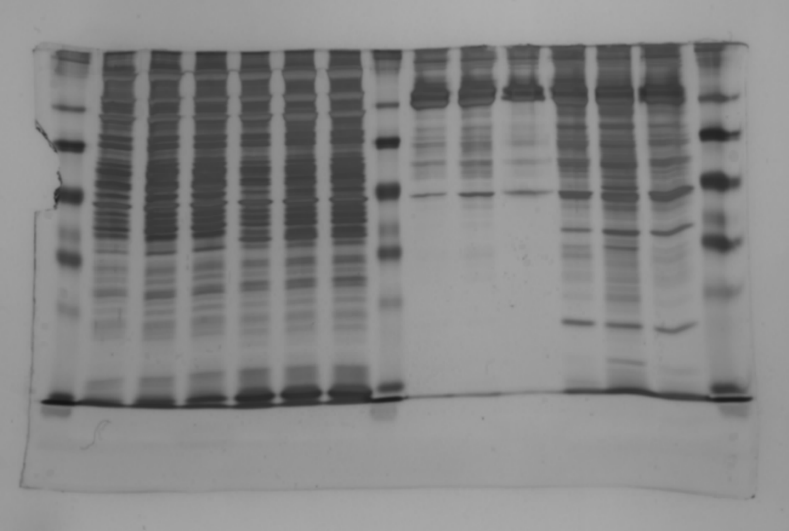

**a**

**b**

**c**

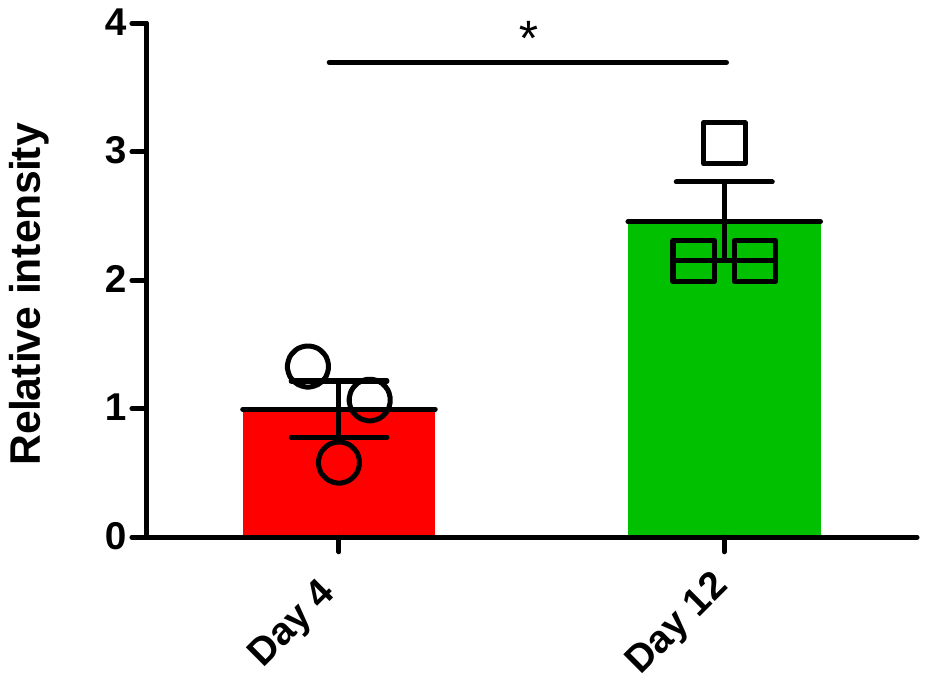

Insoluble fraction

Soluble fraction

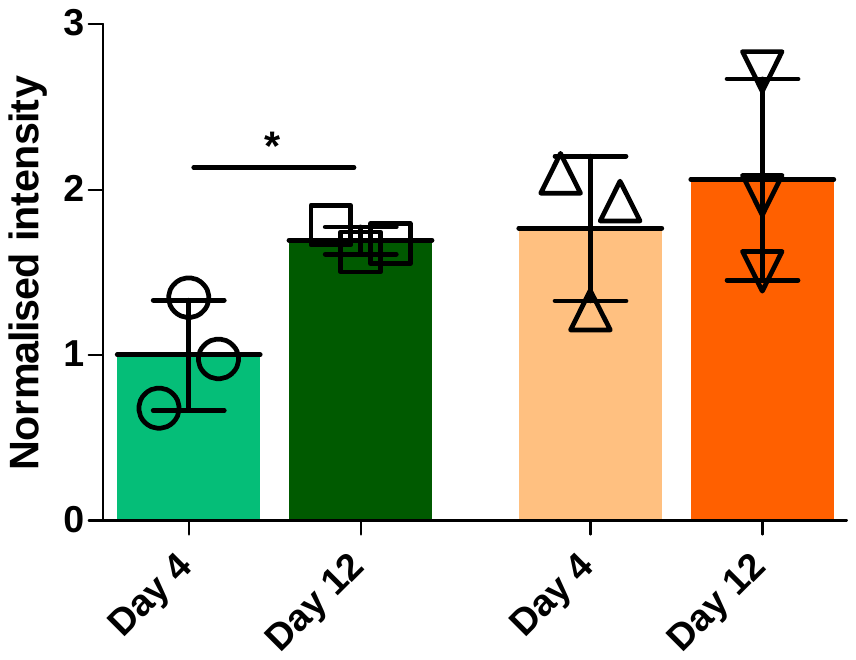

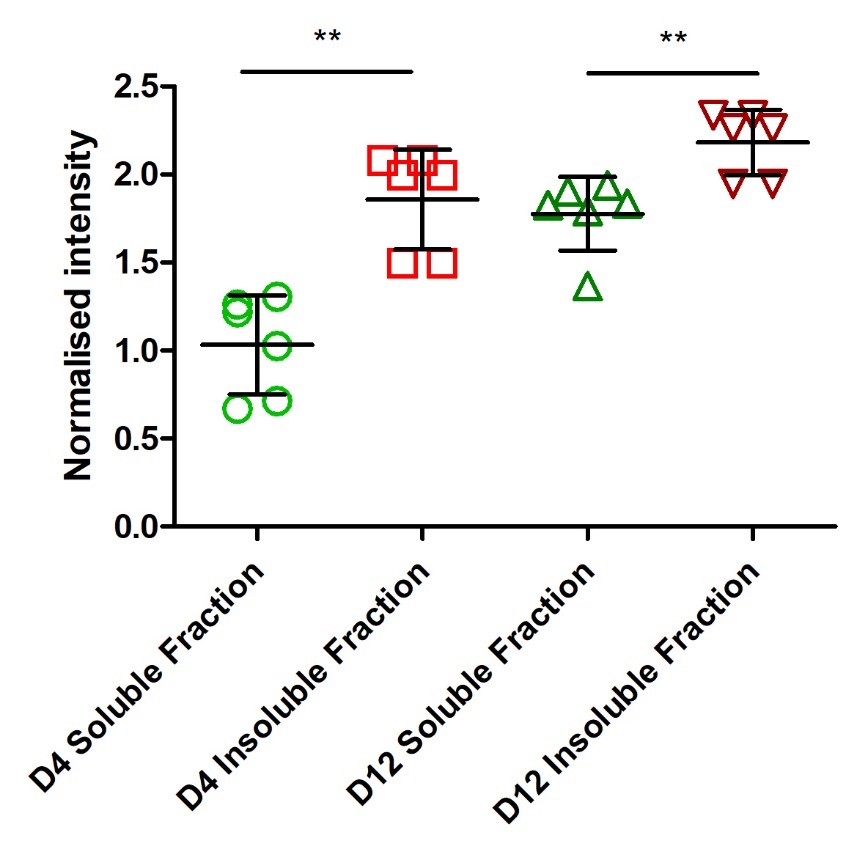

**d**

**Figure S1. Age-dependent accumulation of detergent-insoluble protein and oxidatively damaged protein.** (a) Comparison of detergent-insoluble protein between nematodes aged four days and 12 days relative to detergent-soluble protein (n=three independent cohorts per age). Fractions of soluble and insoluble protein were loaded on SDS-PAGE gel and silver-stained. The red box indicates low molecular-weight detergent-insoluble protein showing a significant age-dependent increase. (b) Detergent-insoluble protein (<49 kDa, inside of the area outlined in red) was quantified using densitometry (ImageJ, NIH, USA). Analysis confirms a statistically significant increase with age between Day 4 and Day 12 (fold change of 2.5±0.5 (mean±SD), p<0.05, paired t-test). (c) PCC was normalized to total protein in soluble fractions (SFs) and insoluble fractions (IFs). Densitometry results of each measurement were normalized to the mean of the intensity measured in the SF of Day 4 animals. In the SFs, there was a 1.8±0.4 fold increase in normalized PCC between Day 4 and Day 12 (mean±SD, p<0.05, t-test). (d) Each repeat of PCC for soluble and insoluble on each day was normalized to the average of Day 4 soluble fraction. The comparison between soluble fraction and insoluble fraction for each Day 4 and Day 12 showed that the insoluble fraction is more oxidized than the soluble fraction overall (p<0.01 for both Day 4 and Day 12, paired t-test).

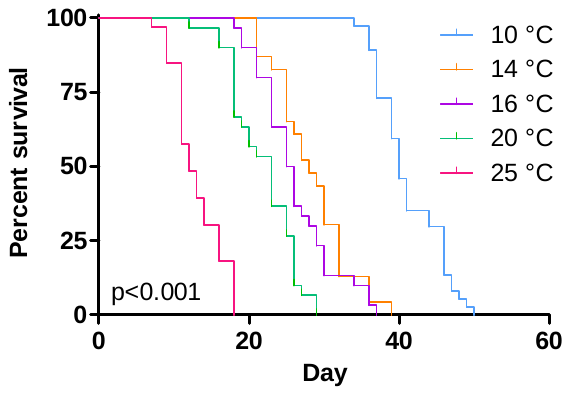

**a**

**b**

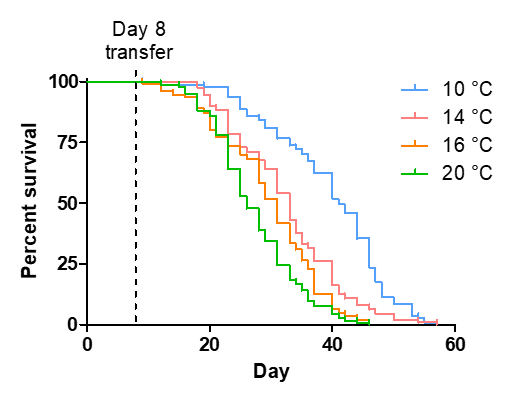

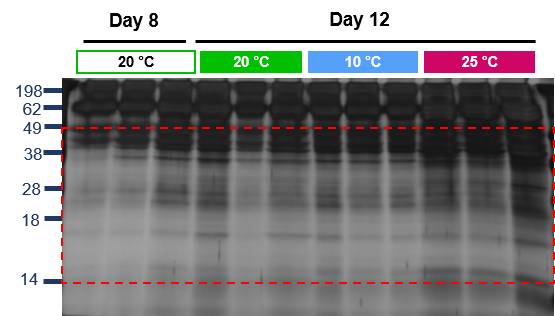

**c**

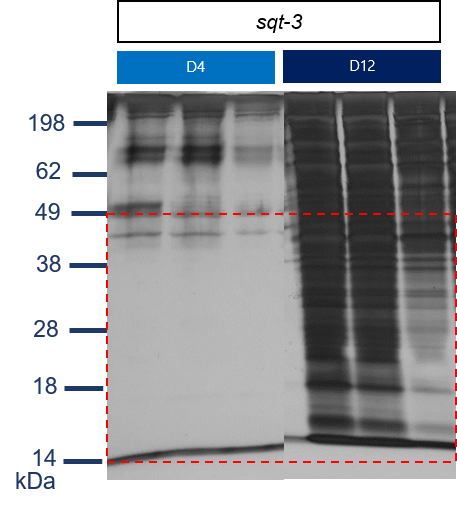

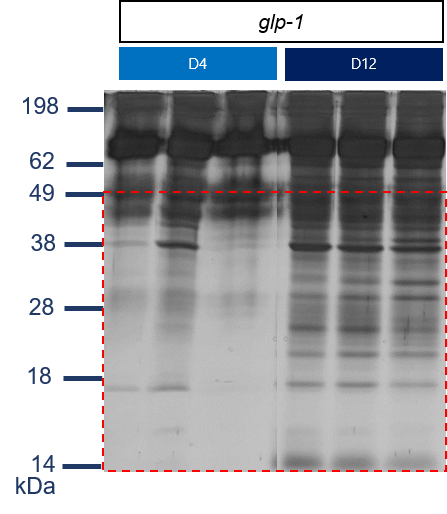

**d**

**Figure S2**. **Insoluble protein accumulate with temperature and with intact oogenesis.** (a) Survival curves of WT *C. elegans* grown at different temperatures throughout life. Temperatures range from 10 to 25 °C and significantly impact lifespan (p<0.001, Log-rank test). (b) The survival curves of WT *C. elegans* maintained at 20 °C until Day 8, then transferred to 10, 14, and 16 °C for the rest of life. Median lifespan: 10 °C, 41 days (n = 141), 14 °C, 33 days (n = 111), 16 °C, 31 days (n = 110), 20 °C, 26 days (n = 142). (c) Gel image for insoluble protein extracted from populations of nematodes raised at 20 °C until Day 8 and transferred for 10, 16, and 25 °C and one group maintained at the same temperature. (d) Gel images for *glp-1* and *sqt-3* mutants fed with bacteria carrying empty vectors (EV), with insoluble protein extracted on both Day 4 and Day 12.

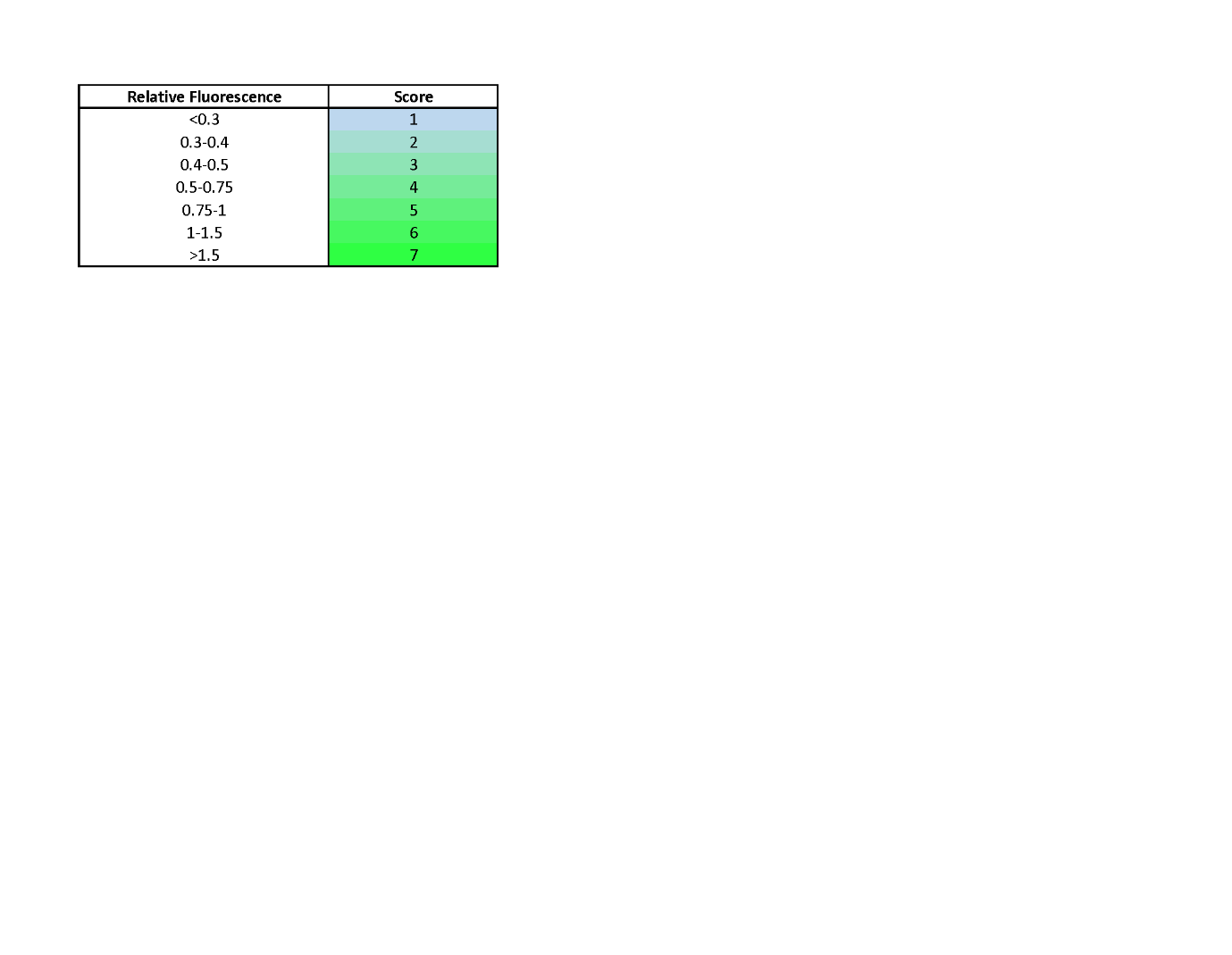

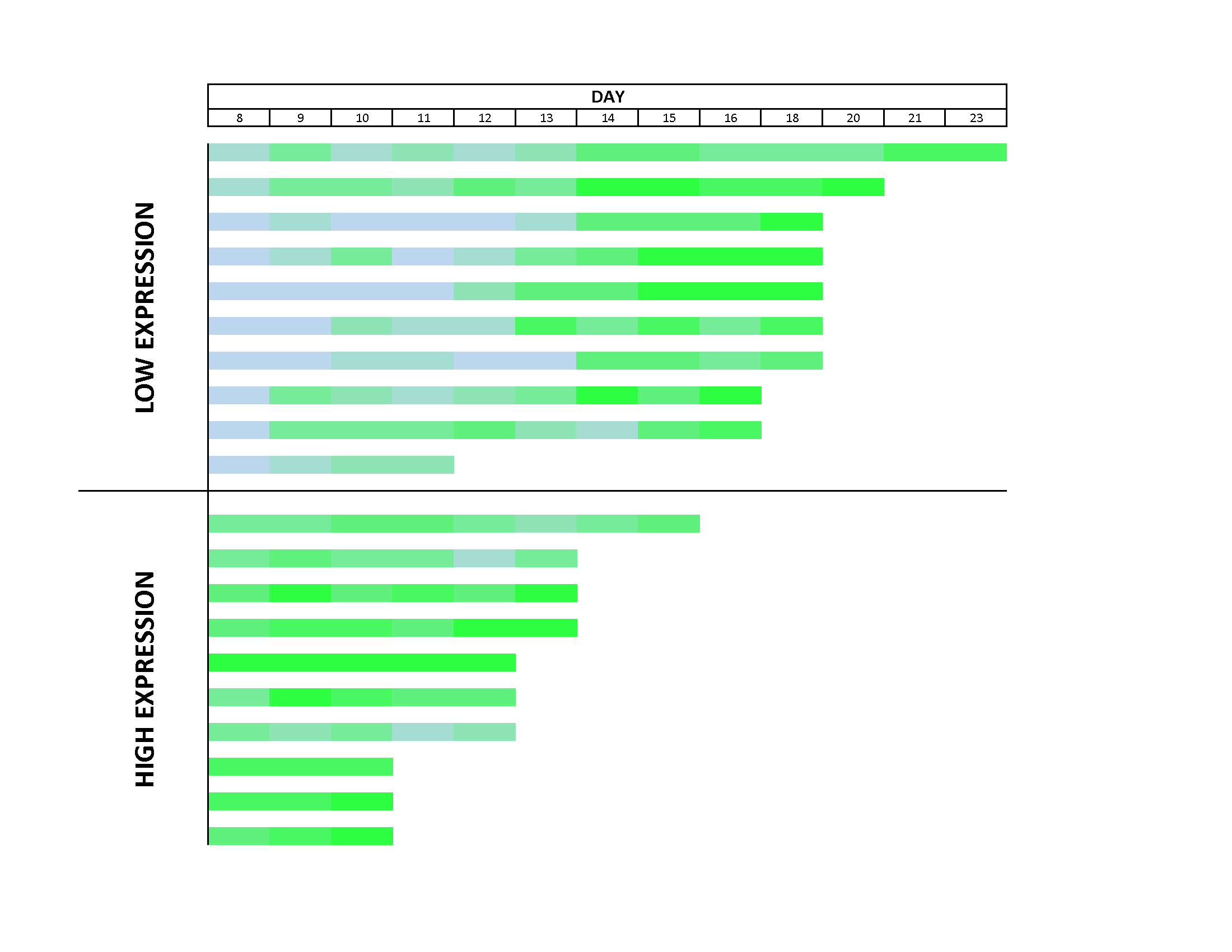

**Figure S3**. ***hsp-16.2* activation increases with age irreversibly.** Heat map showing relative fluorescence for ten individual bright and dim worms maintained at 25 °C and sorted on day 8 of adulthood. Worms were binned into scored groups based on brightness to observe fluorescence dynamics over time. Ends of individual bars indicate the death of the individual.

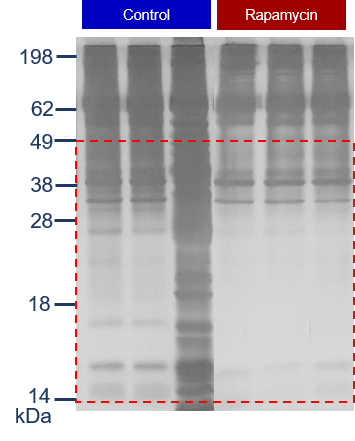

**Figure S4. Comparison of insoluble proteins between control and rapamycin-treated worms.** (**a**) Full gel image for insoluble protein extracted from populations of control and worms maintained on agar plates supplemented with 100 µM rapamycin.

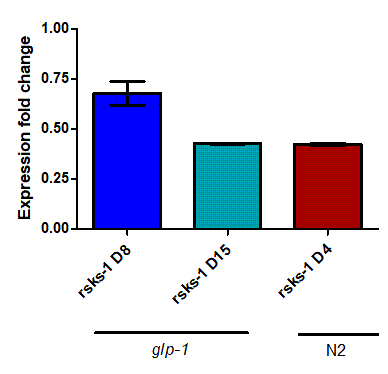

**(iii)**

**(ii)**

**(i)**

**a**

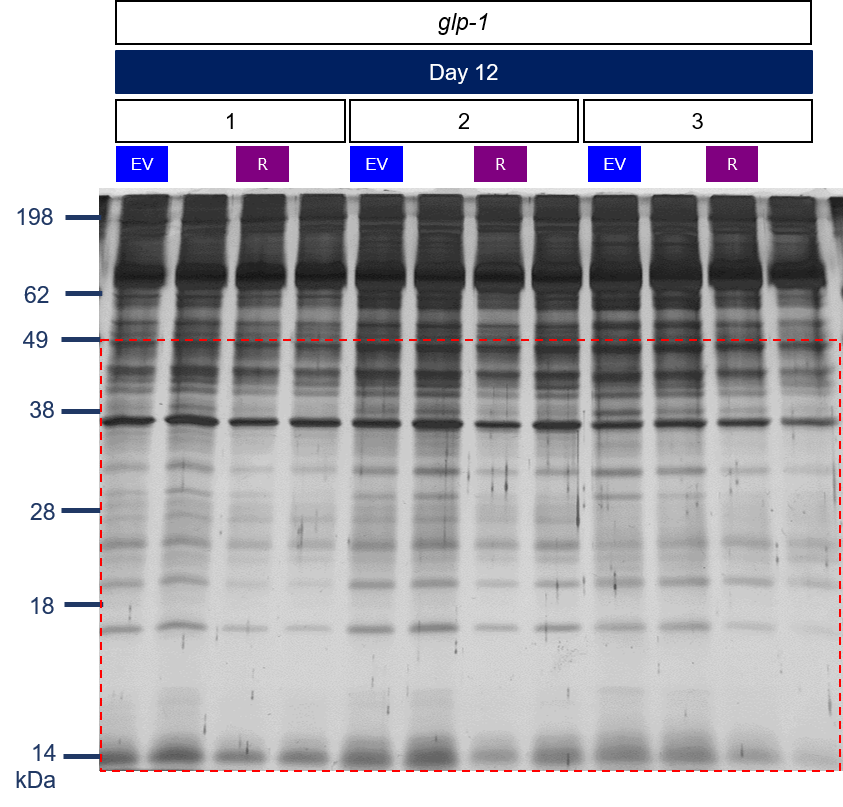

**b**

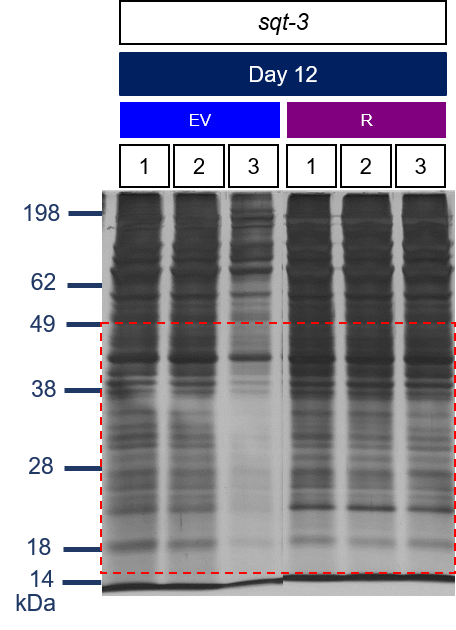

**c**

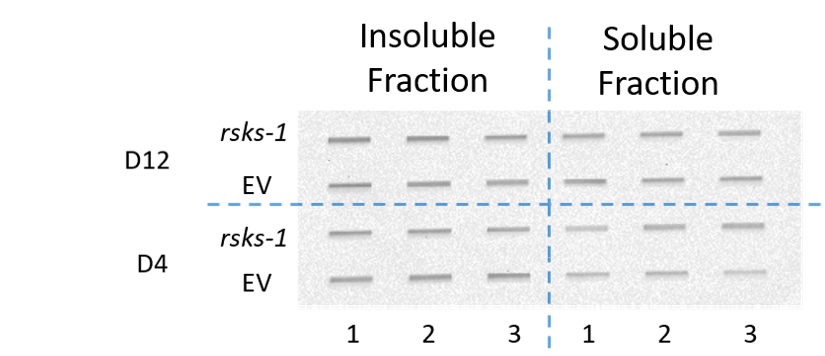

**d**

**Figure S5. Comparison of insoluble protein and protein oxidation in *rsks-1* knockdown animals.** (**a**) Successful knockdown of RNAi conditions shown by qPCR measurement. Measurement of gene expression fold change in the *glp-1* mutants and WT animals (N2) fed with *rsks-1* RNAi. Gene expression fold change measurement shown for *rsks-1* RNAi treatment, (i) Day 8 *glp-1* mutants, (ii) Day 15 *glp-1* mutants, and (iii) Day 4 N2 wild-type animals, normalised to a housekeeping gene, *act-1* and to control (EV). Data represent Mean±SD. (**b-c**) Gel images for insoluble protein extracted from populations of control (EV) and *rsks-1* RNAi bacteria (R) of (**b**) *glp-1* (grouping indicated above lanes) and (**c**) *sqt-3* mutants. (**d**) PCC of soluble and insoluble fractions of animals fed with *rsks-1* RNAi. A full blot image with protein extracted from young (Day 4, D4) and old (Day 12, D12) populations of *glp-1*, control (EV), and fed with *rsks-1* RNAi bacteria. The detergent-soluble and -insoluble proteins were derivatized, transferred to a membrane, and blotted for chemiluminescence detection.

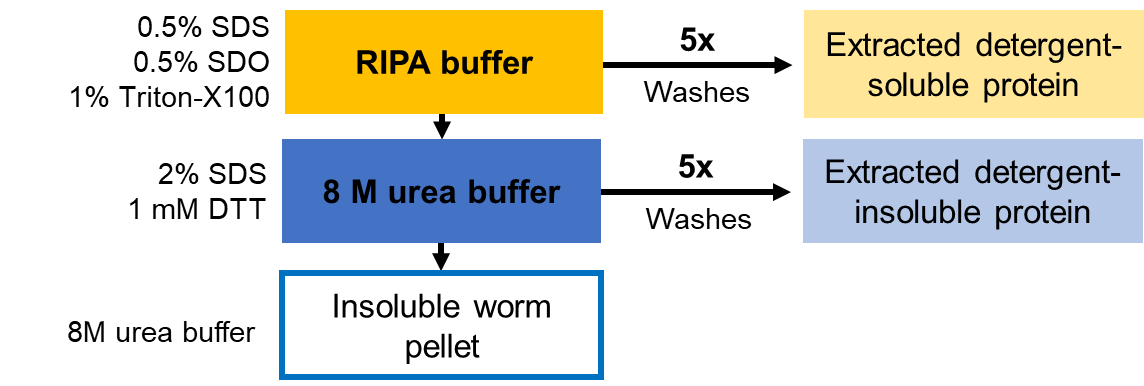

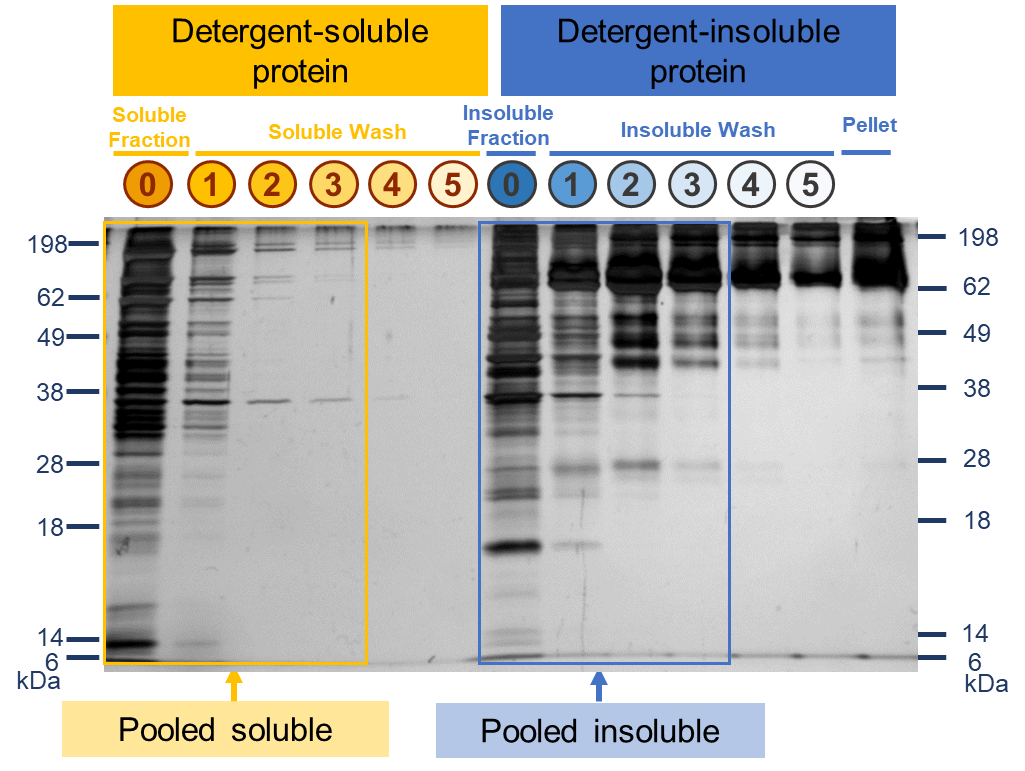

**Figure S6**. **Sequential washing steps for protein extraction.** (**a**) Samples were collected, then detergent-soluble and detergent-insoluble protein were extracted, using sequential washes, and lastly, the remaining pellet was obtained. (**b**) Proteins from each wash step were then loaded on SDS-PAGE, and the quantity of protein extracted for each step was evaluated.

**Table S1. Statistical analysis for the lifespan data using Online Application for Survival Analysis 2, OASIS 2 (Yang et al., 2011)**

|  | | | | **Restricted mean lifespan±S.E.M. (Days)** | **Comparison** | **Change in restricted mean lifespan** | **Log-rank test** | **Survival time at % mortality (Days)** | | | |
| --- | --- | --- | --- | --- | --- | --- | --- | --- | --- | --- | --- |
| **Figure** | **Group** | **Strain** | **n** |  |  |  | **p-value** | **50%** | **p-value** | **90%** | **p-value** |
| **1d** | EV *glp-1* | JK1107 | 106 | 19.37±0.33 | *sqt-3* vs. *glp-1* | 88% | <0.001 | 18 | <0.001 | 22 | 0.55 |
|  | EV *sqt-3* | CB4121 | 119 | 17.07±0.39 |  |  |  | 14 |  | 20 |  |
| **2c** | 20 °C | CL2070 | 105 | 19.84±0.44 | 25 °C vs.20 °C | 70% | <0.001 | 17 | <0.001 | 22 | <0.001 |
|  | 25 °C |  | 109 | 13.79±0.36 |  |  |  | 11 |  | 17 |  |
| **2e** | 1% low expression | CL2070 | 95 | 19.34±0.45 | High vs. low | 81% | <0.001 | 16 | <0.001 | 22 | <0.01 |
|  | 1% high expression |  | 100 | 15.68±0.33 |  |  |  | 14 |  | 16 |  |
| **2f** | 10% low expression | CL2070 | 48 | 20.85±0.7 | High vs. low | 91% | <0.05 | 19 | 0.32 | 24 | 0.46 |
|  | 10% high expression |  | 57 | 18.98±0.52 |  |  |  | 16 |  | 21 |  |
| **3a** | Control | CL2070 | 109 | 13.79±0.36 | Rapamycin vs. Control | 109% | <0.05 | 11 | <0.05 | 17 | 0.78 |
|  | Rapamycin |  | 111 | 14.99±0.37 |  |  |  | 14 |  | 17 |  |
| **4a** | EV (Control) | N2 | 132 | 19.39±0.36 | *rsks-1* RNAi vs. EV | 105% | <0.05 | 16 | 0.09 | 22 | 0.09 |
|  | *rsks-1* RNAi |  | 79 | 20.33±0.61 |  |  |  | 16 |  | 24 |  |
| **4b** | EV (Control) | CB4121 | 119 | 17.07±0.39 | *rsks-1* RNAi vs. EV | 109% | <0.01 | 14 | 0.09 | 20 | <0.05 |
|  | *rsks-1* RNAi |  | 160 | 18.53±0.41 |  |  |  | 14 |  | 22 |  |
| **4c** | EV (Control) | JK1107 | 106 | 19.37±0.33 | *rsks-1* RNAi vs. EV | 95% | 0.09 | 18 | 0.47 | 24 | 0.36 |
|  | *rsks-1* RNAi |  | 98 | 18.49±0.47 |  |  |  | 16 |  | 24 |  |

n.a.: not applicable

**Table S2. Statistical analysis for non-linear curves fitting of the percentages of bright worms over time using GraphPad PRISM**

|  | Control | Rapamycin |
| --- | --- | --- |
| Sigmoidal, 4PL, X is time |  |  |
| Best-fit values |  |  |
| Top | 95.18 | 96.42 |
| Bottom | -15.86 | -11.13 |
| 50% of the population was“bright” | 6.273 | 8.550 |
| HillSlope | 0.2576 | 0.1778 |
| Span | 111.0 | 107.5 |
| Std. Error |  |  |
| Top | 1.462 | 2.288 |
| Bottom | 10.27 | 8.121 |
| 50% of the population was “bright” | 0.4122 | 0.4790 |
| HillSlope | 0.03856 | 0.02659 |
| Span | 10.81 | 9.347 |
| 95% Confidence Intervals |  |  |
| Top | 92.08 to 98.28 | 91.57 to 101.3 |
| Bottom | -37.64 to 5.918 | -28.35 to 6.088 |
| 50% of the population was “bright” | 5.399 to 7.147 | 7.535 to 9.566 |
| HillSlope | 0.1759 to 0.3394 | 0.1215 to 0.2342 |
| Span | 88.12 to 134.0 | 87.73 to 127.4 |
| Goodness of Fit |  |  |
| Degrees of Freedom | 16 | 16 |
| R square | 0.9851 | 0.9852 |
| Absolute Sum of Squares | 305.8 | 356.6 |
| Sy.x | 4.372 | 4.721 |
| Number of points |  |  |
| Analyzed | 20 | 20 |

The non-linear curves were compared using a t-test (mean±S.E.M., df=16).

**Table S3. List of primers used for qPCR**

| **Gene** | **Forward primer sequence (5’-3’)** | **Reverse primer sequence (5’-3’)** |
| --- | --- | --- |
| *rsks-1* | CCGTTTGTGGGATTCACC | TGGCTTTCTCGGGCTCTT |
| *act-1* | GAGCACGGTATCGTCACCAA | TGTGATGCCAGATCTTCTCCAT |
